# Computational characterization of metacognitive ability in subjective decision-making

**DOI:** 10.1101/2025.05.23.655775

**Authors:** Corey R. Plate, Dhruv Govil, Charles Y Zheng, Zoe M. Boundy-Singer, Corey M. Ziemba, Silvia Lopez-Guzman

## Abstract

Metacognition is the process of reflecting on and controlling one’s own thoughts and behaviors. Metacognitive ability is often measured through modeling the relationship between confidence reports and choice behavior in tasks where performance can be objectively measured. Previous work has explored whether metacognitive ability is conserved across different types of tasks, and across different domains such as perception and memory. However, it is unclear whether approaches to the assessment of metacognitive ability that have worked in these contexts can be extended to value-based decision-making where objective accuracy cannot be evaluated. Here, we compare metacognitive ability across different tasks spanning perception and valuation, using Bayesian hierarchical estimation of the parameters of a computational process model of confidence. This model captures metacognitive ability as the uncertainty of an individual’s own decision uncertainty, and can do so in the space of any decision variable, regardless of whether it indexes external features or internal, subjective states. We find that metacognitive ability can be reliably estimated in both objective and subjective decision-making tasks, and is relatively well conserved across tasks, especially within domains and similar confidence reporting paradigms.

## Introduction

Monitoring and reasoning about the quality of our thoughts and behaviors is essential for making better decisions and improving outcomes. This “metacognitive” ability is often studied by prompting participants to assess their confidence in each of a series of decisions. Every decision we make is automatically accompanied by a sense of confidence, which is generally related in a lawful way to the accuracy or reliability of that decision ^1–4^. However, there are large individual differences in the way that people report confidence, likely reflecting broader variation in metacognition and its underlying neural mechanisms across the population ^5–7^. Many studies have explored individual differences in metacognition, with a focus more recently on how metacognitive dysfunction may relate to psychopathology ^8,9^. One major construct of interest is the quality of metacognition, or “metacognitive ability.” This ability is often operationally defined as the degree to which confidence reports reliably reflect the probability of making a correct decision, isolated from overall performance level or the propensity to report high confidence ^10,11^. Thus, studies of metacognitive ability have almost exclusively focused on decisions that have a correct answer, such as memory or cognitive tasks, perceptual judgments, or trivia questions ^12,13^. However, humans also introspect about decisions that have no objectively correct or incorrect answer ^14^. For example, when making a choice about what car to purchase, a decision-maker may not be guided by which car is objectively better. Instead, they will rely on an internally constructed subjective value that should lead them to choose the car that is best for them. A feeling of confidence will accompany that type of choice as well, but metacognitive ability in such preference-based decisions remains under-characterized and largely unexplored ^15^. Here, we address this gap by extending and combining computational frameworks for value-based decisions and confidence generation, allowing us to reliably assess metacognitive ability in subjective decision-making.

Research on metacognition and confidence has grown significantly over the last decade ^16^. However, only a handful of studies have focused on value-based as opposed to perceptual or memory-guided decisions. Nonetheless, our work has been propelled by recent efforts at investigating how confidence can emerge from value-related processes ^14,15,17–19^. Most value-based decision models rest on the assumption that the subjective value of different options are first computed and subsequently compared to implement a choice. In one of the first demonstrations of the relation between confidence and value, De Martino et al. showed that confidence in a choice between two food items increased with the magnitude of the subjective value difference between them ^14^. They directly inferred the subjective value of each food item by asking participants to report their willingness-to-pay in a bidding task ^20^. Strikingly, decisions made with high confidence were more strongly associated with the subjective value difference between choice options, as if the participants could assess the reliability of their internal decision variable and report high confidence when their decision was more likely to correspond to their preferences expressed in the bidding task ^14^. Further studies suggest that confidence is influenced by noise in the decision process of comparing values, rather than in an earlier encoding of value representations ^14,21^. This supports the interpretation that confidence reflects a cognitive appraisal ^22^ of the quality of the decision itself rather than of the quality of the stimuli or option values, despite evidence that confidence information may be “automatically” decodable from areas like ventromedial prefrontal cortex that also represent decision primitives such as value, probability, or pleasantness ^23^. Finally, participants show some variation in the difference in association between confidence and the subjective decision variable, an indication that metacognitive ability in these value-based tasks may differ between individuals ^15^.

That confidence behavior reflects a metacognitive appraisal of decision quality and thus may be informative about an individual’s ability to introspect, has become a topic of broad interest ^24,25^. However, quantitatively assessing an individual’s metacognitive ability from choice-confidence data is challenging. Many straightforward measures can be confounded with overall performance level or the propensity to report higher confidence ^10,11,26^. Models rooted in signal detection theory ^27^ have allowed for some isolation of measures of sensitivity and bias in confidence reporting ^11,26,28–32^. Importantly, these models have driven an increasing number of studies that relate individually estimated measures of metacognitive ability to differences in other behaviors, neuropsychiatric conditions, or variation in psychopathology ^8,33–37^. These *descriptive* models measure metacognitive ability as the extent to which confidence differentiates between correct and incorrect trials ^11,28,29,38^. However, there is increasing recognition that confidence reports generally do not reflect a participant’s estimate of making a correct decision (the so-called “Bayesian confidence hypothesis”) ^1^, even for simple perceptual decisions ^39–44^. Thus, differentiating correct or incorrect trials may not be the most appropriate rubric for judging metacognitive ability. Instead, confidence reports seem more consistent with an estimate of choice consistency or decision reliability ^42,45,46^. This idea has recently been instantiated in a *process* model of confidence generation that treats confidence as a noisy decision reliability estimate (the CASANDRE model) ^46^. In this model, metacognitive ability is limited by the uncertainty in the estimated uncertainty in the internal decision variable, a quantity thus designated “meta-uncertainty.” Other recently proposed models can capture metacognitive ability with similar parameters (referred to as meta-, metacognitive, or confidence noise) ^47–50^. Such an approach can facilitate the study of the quality of metacognition even for subjective decisions.

There are initial suggestions that estimates of metacognitive ability from a specific task may generalize and indicate something broader about an individual’s cognitive capacities. Metacognitive ability is at least somewhat conserved across tasks that target different domains of cognition such as perceptual decision-making, mathematical problem solving, episodic memory, and general knowledge or trivia ^13,46,51–53^. This has important implications, as it provides support for the internal validity of task-based measures of metacognitive ability and their potential use for capturing clinically relevant features of metacognition in psychiatric populations. However, this previous work exclusively focuses on decisions where there is a correct or incorrect answer, and has not extended to the type of decisions most often affected across psychopathology ^54^. Altered value-based decision-making, such as excessive approach or avoidance of risk and choosing immediate rewards at the expense of future outcomes, are pervasive behaviors across psychiatric disorders ^55,56^ such as addiction ^57,58^, gambling disorder ^59^, anxiety ^60^, or ADHD ^61^. These behaviors are typically captured in tasks where participants are asked to choose what they prefer instead of what is correct. Whether individual differences in metacognitive ability can be reliably modeled across these types of subjective decisions remains an open question.

Here, we leverage two technical innovations to address whether metacognitive ability can be estimated using choice-confidence data from preference-based decision-making tasks, whether it is conserved across different value-based decision-making tasks, and whether it generalizes across objective and subjective decision-making. First, we combine utility models and the CASANDRE model ^46^ to explain confidence reporting behavior in two preference-based tasks: a delay discounting task and a risk and ambiguity task. This procedure allowed us to computationally characterize metacognitive ability during subjective decision-making for the first time. Second, we developed a Bayesian hierarchical estimation routine for optimization of the CASANDRE model and for estimation of cross-task correlations. These methods allowed us to reliably infer model parameters from data with many participants but few trials per participant, as well as the dependency between parameters across tasks. We find that metacognitive ability is indeed a trait-like construct that generalizes strongly in single participants across different value-preference tasks, although less so between tasks that differ across domains of decision-making and the type of confidence report. Our methods and results further open up the field of metacognitive research into subjective decision-making – a topic of considerable interest in the field of metacognition ^24,62,63^ – and also bolster previous claims that metacognitive ability is an internally and externally valid construct with implications for many domains of psychological and psychiatric research.

## Results

To evaluate metacognitive ability in subjective decision-making, we used two well-established tasks for which quantitative models of behavior have been widely implemented: a delay discounting task and a risk and ambiguity task. Behavior in these two tasks reflects distinct and dissociable preferences ^64,65^. Choice behavior in the delay discounting task reflects an individual’s willingness to wait for rewards. Choice behavior in the risk and ambiguity task reflects an individual’s preference for probabilistic outcomes when the probabilities are fully known, or risk tolerance, as well as their preference for probabilistic outcomes when the probabilities are not fully known, or ambiguity tolerance. Each trial of the delay discounting task involves a binary choice between a smaller-but-sooner and a larger-but-later monetary reward, with delays and amounts varying trial-by-trial (Fig. 1a). After each choice, participants were asked to rate their confidence that their choice reflected their true preference on a scale from 1 to 4. Similarly, each trial of the risk and ambiguity task involved a choice between a certain small monetary reward and a lottery that could pay a larger amount or nothing (Fig. 1b). The reward probabilities and amounts of the lottery varied trial-by-trial. In risky trials, all the information about probabilities was shown, but in ambiguity trials, part of this probability information was masked. As in the delay discounting task, choices in the risk and ambiguity task were followed by a confidence rating. We incentivized participants to choose according to their preferences by instructing them that any of their choices could be chosen at the end of the experiment and that its outcome would reflect their bonus payment.

**Figure 1.**
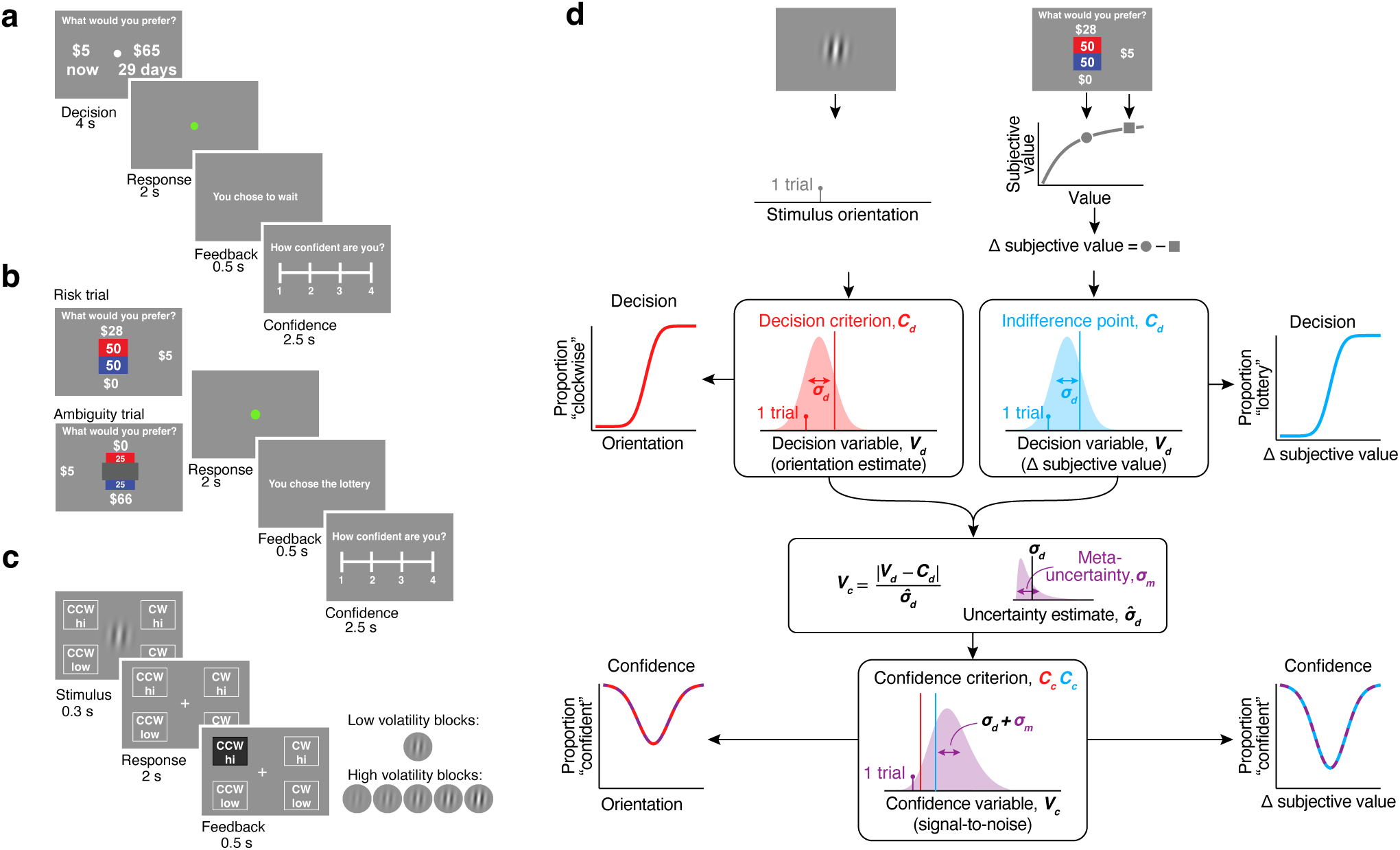
A process model of decision confidence can conceptually account for the quality of metacognition across perceptual and subjective decision-making tasks. **a**, Experimental design for the delay discounting task. **b**, Experimental design for the risk and ambiguity task. **c**, Experimental design for the orientation discrimination task. Participants performed two blocks where the volatility of stimulus uncertainty was low and only one contrast was presented, and two blocks where the volatility of stimulus uncertainty was high and five different contrast levels were randomly interleaved. **d**, Schematic of the hierarchical CASANDRE modeling framework and illustration of the hypothesis of a shared metacognitive processing stage across objective and subjective decisions.

Generative models of confidence have not previously been applied to subjective decision-making behavior, but can be used to effectively explain confidence reports in perceptual decisions ^46–50,66^. To compare confidence reporting behavior in subjective decision-making with more commonly studied perceptual confidence behavior, we employed a simple orientation discrimination task. Participants judged whether a grating stimulus was tilted clockwise or counterclockwise from vertical (Fig. 1c). However, unlike many studies of perceptual confidence, our study varied both the stimulus strength (the magnitude of the tilt from vertical) and the stimulus uncertainty (the contrast of the grating) on each trial. This complexity mirrors that of the value-based tasks and necessitates a model for confidence that can accommodate diverse levels of evidence strength. We used a confidence reporting procedure that differed from that in the value-based tasks and from much of perceptual confidence literature ^11,67–69^. Participants communicated both their binary orientation and confidence judgments with a single button press from four available options (Fig. 1c). To encourage task engagement similar to the preference-based tasks and incentivize reasonable confidence reporting behavior, the potential reward on each trial depended on both the accuracy of the perceptual judgment and the level of confidence reported (see methods) ^67^. Participants performed two versions of the task, one where the trial-by-trial volatility of stimulus uncertainty was low (one contrast level) or high (five interleaved contrast levels).

### Confidence in subjective decisions as a noisy decision reliability estimate

The richness of value-based decision-making data poses both conceptual and practical challenges for modeling confidence reporting behavior ^14^. Confidence is often conceived as a participant’s estimate of their probability of making a correct decision ^1,2,70^, a framework that clearly does not apply to preferential decisions. Instead, we conceive of confidence as an estimate of decision reliability that reflects a participant’s impression of their probability of self-consistency. In other words, confidence reports indicate how likely a person would be to make the same decision when presented with the options again ^42,45,46^. However, this estimate can be noisy, and the magnitude of this noise will determine a participant’s metacognitive ability. Standard methods for assessing metacognitive ability rely on defining it as the degree to which confidence reflects accuracy, and also, practically, on keeping the strength of evidence constant across many trials ^11,29,71^. One of the most difficult aspects of assessing metacognitive ability is to dissociate sensitivity in confidence reporting behavior from sensitivity in decision behavior ^10,26,28^. This can be largely accomplished by expressing the quality of confidence reports (*meta*-*d^↑^*) in the same units used to assess decision sensitivity (*d^↑^*), and taking the ratio to obtain a measure of “metacognitive efficiency” (*meta*-*d^↑^/d^↑^* or M-ratio) ^11,29^. However, this approach requires tasks where decision evidence is held or assumed to be roughly constant, allowing for repeated trials that will generate the same level of accuracy. This is difficult to attain in preference-based choices where the decision variable is internal to the decision-maker because it results from a process of idiosyncratic option valuation and comparison.

Instead of standard approaches for quantifying metacognitive ability, we turned to a generative process model that captures confidence as a noisy decision reliability estimate (CASANDRE) ^46^. This model has been applied to behavior in diverse perceptual decision-making and confidence tasks in both human and nonhuman primate participants ^25,46,67,72^. It posits a two-stage process for decision-making and confidence estimation rooted in traditional signal detection theory ^46^. In the first stage, a noisy decision variable is compared against an internal decision criterion to generate a binary choice (Fig. 1d). In the case of orientation discrimination, this decision variable is a direct internal estimate of the external stimulus orientation (Fig. 1d, top left). In the second stage, the distance between the decision variable and the decision criterion is normalized by an estimate of the uncertainty (or noisiness) of the decision variable. The resulting confidence variable expresses an estimate of decision reliability (as a signal-to-noise ratio) and is compared to a confidence criterion to generate a binary confidence report (or multiple criteria to produce a confidence rating; Fig. 1d, bottom). The critical innovation of the CASANDRE model is that it assumes the uncertainty estimate itself is uncertain ^46^. This uncertainty about uncertainty is referred to as “meta-uncertainty” and captures the accuracy with which confidence reports reflect decision reliability, a measure of metacognitive ability. In perceptual and other objective decision-making tasks, meta-uncertainty shows some hallmarks of a trait-like construct, but can also be experimentally manipulated and potentially mitigated with learning ^46,67^.

We propose that confidence reports in subjective decisions are generated through a process analogous to the CASANDRE account of perceptual decision-making ^46^ (Fig. 1d, right). However, in the case of subjective decision-making, the first-stage decision variable is not a direct reflection of an external quantity (e.g., orientation) but an internal subjective quantity that needs to be inferred from choice behavior in the tasks. To do this, we employed a discounted utility model ^65^ and an expected utility model ^73,74^ in the delay discounting task and risk and ambiguity task. In these utility models, the objective variables of the decision (delay and amount for the delay discounting task, and probability, ambiguity, and amount for the risk and ambiguity task) are computed and transformed into a utility or subjective value for each option. Subjective values are then compared and the resulting subjective value difference is passed through a stochastic decision rule ^75,76^ leading to a binary choice. The trial-by-trial subjective value difference estimated from the utility models was thus our first-stage decision variable (Fig. 1d, top right).

We hypothesized that if metacognitive ability reflects a general, trait-like process, possibly limited by shared neural mechanisms, then our estimate of metacognitive ability derived from the CASANDRE model should generalize across decision-making tasks both within and across the domains of perceptual and value-based decision-making. Previous work has established some degree of internal validity for metacognitive ability, that is, consistency across tasks for measures of confidence efficiency ^12,13,53^. While these studies have focused on perception, memory, and general knowledge tasks that all have objectively correct or incorrect responses, ours is the first study to our knowledge that attempts to computationally characterize metacognitive ability for subjective decision-making tasks as well.

### Predicting confidence reporting behavior across both perceptual and value-based decisions

We recruited 128 participants (52 female, ages 18-55) from Amazon Mechanical Turk using CloudResearch’s MTurk Toolbox. Participants were asked to complete online versions of all three tasks (Fig. 1 a-c). We used monetary incentives to encourage participants to choose according to their true preference in the subjective decision-making tasks and to accurately determine orientation in the perceptual tasks. We accomplished this by randomly selecting a single trial from any of the tasks after all were completed and adding the outcome of the participant’s choice in that trial as a bonus to their payment. Given that any trial had the same probability of getting chosen at the end, participants were encouraged to avoid missing trials or responding randomly. We excluded participants online based on failure to achieve benchmark performance, and post-hoc based on whether their confidence behavior failed to show sufficient variation for meaningful model fitting (see Methods).

Participants exhibited systematic behavior in all three tasks that was well described by the CASANDRE model ^46^. Figure 2 illustrates data and model predictions from 2 example participants with diverse behavioral profiles. For the low volatility perceptual decision-making task, participants 1 and 2 exhibited similar decision sensitivity (as shown by the slope of the psychometric functions; Fig. 2ad, top), but very different levels of confidence in their decisions (Fig. 2ad, bottom). To examine behavior in the delay discounting task, we first fit a discounted utility model ^65^ to a participant’s choices between delayed and immediate offers. We then grouped trials according to similar subjective value differences between the two options (Fig. 2be, top). Computing the proportion of responses in which the participant chose the delayed option in these groups results in a psychometric function whose slope expresses the participant’s decision sensitivity. Note that although each participant saw an identical set of trials, the distribution of subjective value differences is idiosyncratic because it results from each individual’s distinct preferences for delayed rewards. Similarly, for the risk and ambiguity task, we fit an expected utility model ^73,74^, and then constructed psychometric functions by examining the proportion of lottery choices as a function of the subjective value difference between the risky and certain options (Fig. 2cf, top). Here again, different preferences for risk and ambiguity result in different distributions of subjective value difference across individuals. For both subjective decision tasks, the choice proportion approaches 0.5 around a subjective value difference equal to 0, referred to as the indifference point. This point of greatest decision uncertainty is also where confidence reports are lowest across tasks and participants (Fig. 2bcef, bottom). This is similar to the point of highest decision uncertainty for the perceptual decision-making task, corresponding to the internal decision criterion (which can differ from the externally enforced criterion of zero degrees of stimulus orientation; Fig. 2ab).

**Figure 2.**
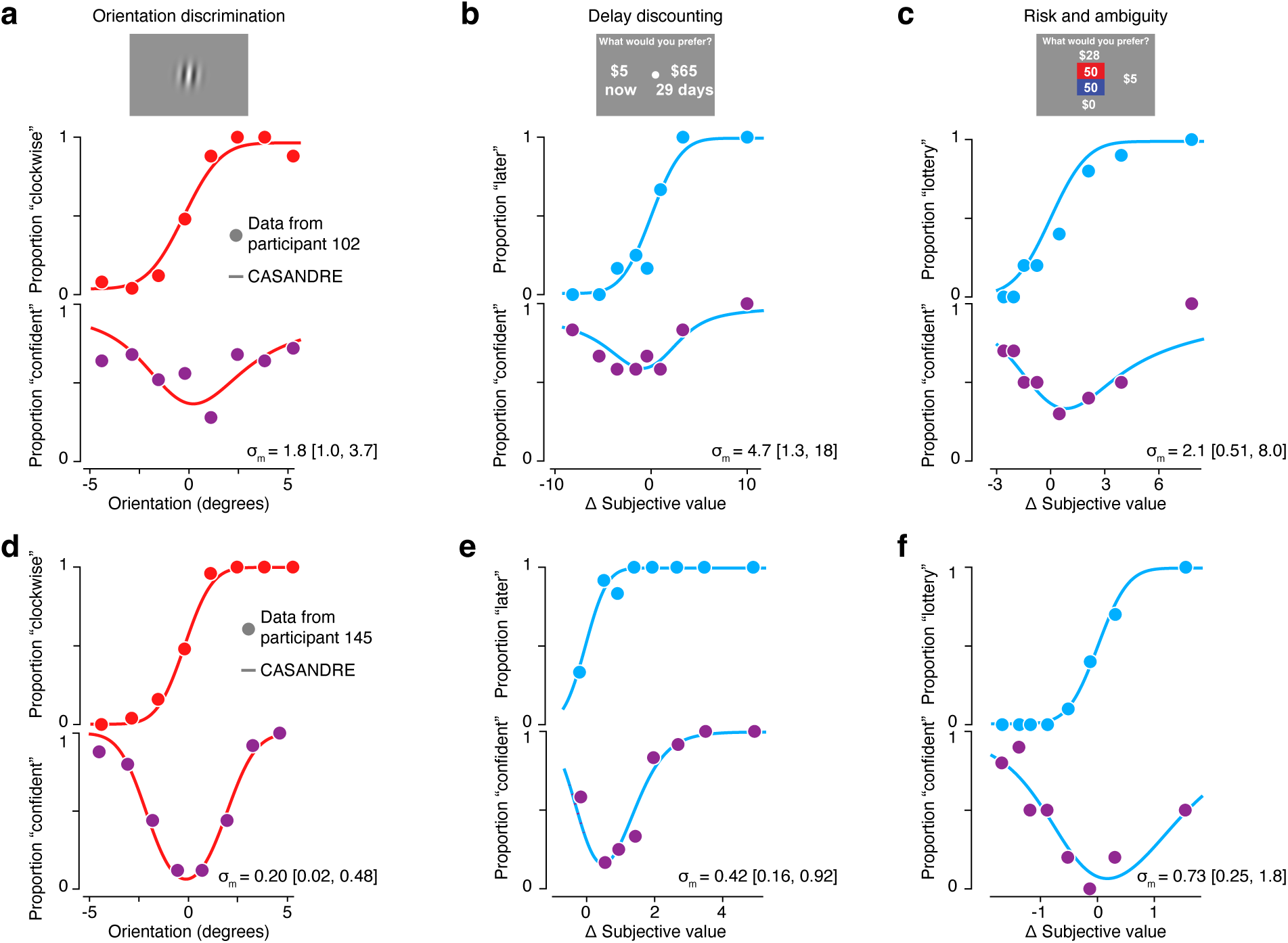
Example behavior across perceptual and value-based decisions. Each panel depicts the psychometric function at the top and confidence function at the bottom for a particular task. All model fits are from the CASANDRE model optimized with Bayesian hierarchical methods. The x-axes for the delay discounting and risk and ambiguity tasks are derived individually from utility modeling. **a-c**, Psychometric and confidence functions for participant 102 who showed generally high levels of meta-uncertainty across three tasks. The geometric mean meta-uncertainty (*ω_m_*) and 95% confidence intervals estimated from each task are shown at the bottom of each panel. Data and model fits are shown for **a**, low volatility blocks from the orientation discrimination task, **b**, the delay discounting task, and **c**, the risk and ambiguity task. **d-f**, Same as **a-c**, but for data collected from participant 145 who showed generally low levels of meta-uncertainty.

The sensitivity of confidence reporting behavior can be assessed through the estimated meta-uncertainty from the CASANDRE model, just as decision sensitivity can be inferred from the slope of the psychometric function. However, unlike decision sensitivity, the value of meta-uncertainty cannot be easily qualitatively discerned from inspecting the psychometric or confidence functions ^46^. Instead, we must fit a model to have the power to assess metacognitive ability. Higher values of meta-uncertainty indicate greater noise in the transformation of an internal decision variable into a confidence variable and are indicative of lower metacognitive ability. Our two example participants exhibited consistently different levels of meta-uncertainty across tasks. One participant’s behavior yielded higher meta-uncertainty across all three tasks (Fig. 2a-c) when compared with the other participant’s behavior (Fig. 2d-f), suggesting they may have relatively lower metacognitive ability. We wondered whether the pattern suggested by these examples would generalize across a large population of participants, providing evidence for the proposal that metacognitive ability represents a stable, trait-like, construct across individuals and different tasks, even in the domain of subjective decision-making.

### Hierarchical Bayesian estimation of metacognitive ability

As has been previously noted, the choice of parameter estimation method for fitting computational models to behavioral task datasets is critical, particularly when dealing with limited numbers of trials ^77–79^. Thus, to reduce parameter estimation noise ^80^ we employed a Bayesian estimation approach with entropy-maximizing and, when available, empirical priors to estimate the parameters of the utility models (see Methods). Using Bayesian estimation for the parameters of the CASANDRE model is substantially more complicated, as the confidence variable is modeled as a ratio distribution with no closed form solution ^46^. Previous work has estimated the parameters of the CASANDRE model through the maximization of likelihood, but we were able to develop a procedure for using Bayesian hierarchical modeling to estimate the model parameters (see Methods and Supplementary Material). Individual and group-level parameters were estimated using Markov Chain Monte Carlo sampling in Stan. We used trace plots and diagnostic statistics (*R*^^^ and ESS) to assess model fit (Supp. Fig. 6a-d). The model adequately explored the posterior as assessed by energy plot and rank plots (Supp. Fig. 6ef), and provided a good qualitative match to different participant’s choice and confidence reporting behavior (Fig. 2).

We further considered how best to measure the generalization of metacognitive ability across different decision-making tasks. We first attempted to use existing packages for Bayesian hierarchical estimation of metacognitive efficiency (M-ratio) and its correlation across tasks ^71^. However, after applying this model to our data we found that the sampled parameter values failed to converge and so cannot provide reliable estimates (Supp. Fig. 3). This is not surprising as the structure of subjective decision-making tasks is ill-suited for the application of the *meta → d^↑^* framework ^11,71^. Instead, we proceeded to consider different procedures for evaluating cross-task correlation in meta-uncertainty. One approach is to separately fit the CASANDRE model to data from each task, obtain point estimates of the meta-uncertainty parameter from each subject, and then to compute the correlation of these estimates across different pairs of tasks. Alternatively, we can jointly estimate the CASANDRE parameters from different tasks and simultaneously estimate their correlation across tasks as a free parameter. This approach necessitates modeling the correlation of two random variables (here, meta-uncertainty) with arbitrary (non-gaussian) marginal distributions, which can be done through the use of a copula (see Methods). Copulas have been used for this purpose in many different fields from engineering and finance to neuroscience ^81^. Critically, so as not to bias the copula correlation parameter estimate (*ε*), we established its prior distribution to be uniform between -1 and 1.

Before applying these methods to our data, we first verified through simulation that our joint Bayesian hierarchical modeling approach yielded accurate and reliable estimates of both the value and dependency of meta-uncertainty across tasks. We simulated choice-confidence data in the two subjective decision-making tasks from populations of participants with CASANDRE model parameters randomly drawn from their entropy-maximizing priors (Supp. Fig. 2). Meta-uncertainty values for each modeled participant were drawn from a bivariate prior for which we varied the level of dependency across 21 correlation values ranging from 0 to 1. We then analyzed each synthetic population of participants either through “separate” Bayesian hierarchical modeling of each task, or through “joint” copula-based Bayesian hierarchical modeling of both tasks (see Correlation recovery section in Methods). Each run of the model generates meta-uncertainty estimates for each participant of the synthetic population in both tasks, and we generally found better recovery of the true meta-uncertainty values from both tasks when estimated with the joint model (Fig. 3a; for delay discounting task, p = 0.004, W = 198; for risk and ambiguity task, p < 0.001, W = 231, Wilcoxon signed rank test). This was generally true regardless of the strength of the ground truth correlation between meta-uncertainty values in the two tasks. The difference in estimation error between “separate” and “joint” modeling and the generative correlation was unrelated for the risk and ambiguity task (*ε* = -0.12, p = 0.61, Spearman correlation), and only weakly related for the delay discounting task (*ε* = -0.4, p = 0.07). Further, we found that joint modeling with copula-based estimation of the correlation between meta-uncertainty values in both tasks led to relatively accurate recovery of the underlying generative correlation value (Fig. 3b). In contrast, separate estimation and correlation of meta-uncertainty values in the two tasks led to severe and systematic underestimation of the true correlation in participants differs across comparisons because some participants passed inclusion criteria for only some of the tasks.

**Figure 3.**
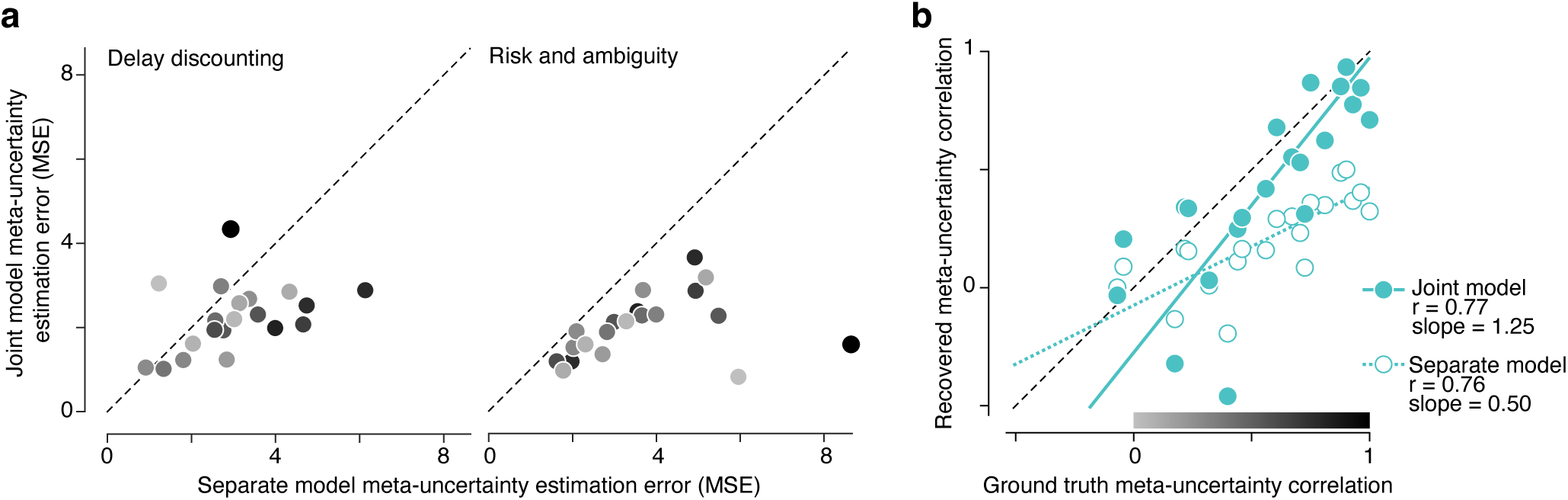
Parameter recovery using copula-based Bayesian hierarchical modeling. **a**, Accuracy of recovered meta-uncertainty estimated from simulated data in the delay discounting (left) and risk and ambiguity (right) tasks. Each symbol represents the mean squared error (MSE) from the ground truth meta-uncertainty values averaged across 105 simulated participants in each synthetic dataset. Separate model fits used data from only one task to estimate meta-uncertainty. Joint model fits used data from both tasks and estimated meta-uncertainty and its cross-task dependency using a copula. Symbols in both panels are shaded according to the color map in **b**, corresponding to the strength of the ground truth correlation. Weak cross-task meta-uncertainty correlations are shaded light gray and strong correlations are shaded black. **b**, Recovery of the cross-task correlation is more accurate with copula-based Bayesian hierarchical modeling. The copula Pearson correlation of the ground truth meta-uncertainty across 105 simulated participants is plotted against the recovered correlation, computed in two different ways. For the joint model (filled symbols), the *ε* value is directly estimated using a copula. For the separate model (open symbols), point estimates of meta-uncertainty in the two tasks are correlated. The colormap at the bottom of the figure serves as a legend for the shading of symbols in **a**.

### Metacognitive ability is conserved across subjective decision-making tasks

As with our modeling of single tasks, we found that fitting the joint CASANDRE model across pairs of tasks similarly passed all model quality checks (Supp. Fig. 6-7). When we applied the model to datasets from the delay discounting and risk and ambiguity tasks, we found that meta-uncertainty was strongly correlated across these two subjective decision-making tasks (Fig. 4a; *ρ* = 0.58, 95% CI = [0.29 *→* 0.8], Bayes factor > 100, decisive evidence). To ensure that this corelation effectively isolated the relationship between metacognitive ability in the two tasks, we examined other decision-making factors that could also be related. This is especially important in this case, as unlike previous work, here we estimate metacognitive ability by inferring an internally generated decision variable. First, we note that overall choice behavior across the two tasks was fairly distinct. We found no significant correlation between the proportion of trials where the lottery was chosen in the risk and ambiguity task and the proportion of trials where the immediate option was chosen in the delay discounting task (*ρ* = 0.16, *p* = 0.1; see Supplement). Still, we wondered whether the correlation in meta-uncertainty across the two subjective decision-making tasks could be explained by shared variance in decision sensitivity, which has previously been found to be correlated in these types of tasks ^82^. We used generalized linear models to predict meta-uncertainty in one task from decision sensitivity estimated from both tasks, and found little evidence that any relationship in decision sensitivity across tasks could be contributing to the conservation of metacognitive ability (see Supplement). Decision sensitivity values across participants explained essentially no variance in the level of meta-uncertainty estimated from either the risk and ambiguity task (*R*^2^ = 0.01) or the delay discounting task (*R*^2^ = *→*0.003). These analyses all indicate that metacognitive ability generalizes across diverse preference-based tasks, and further establishes that our approach toward characterizing metacognitive ability in subjective decision-making provides a relatively stable estimate of a trait-like, individual construct.

**Figure 4.**
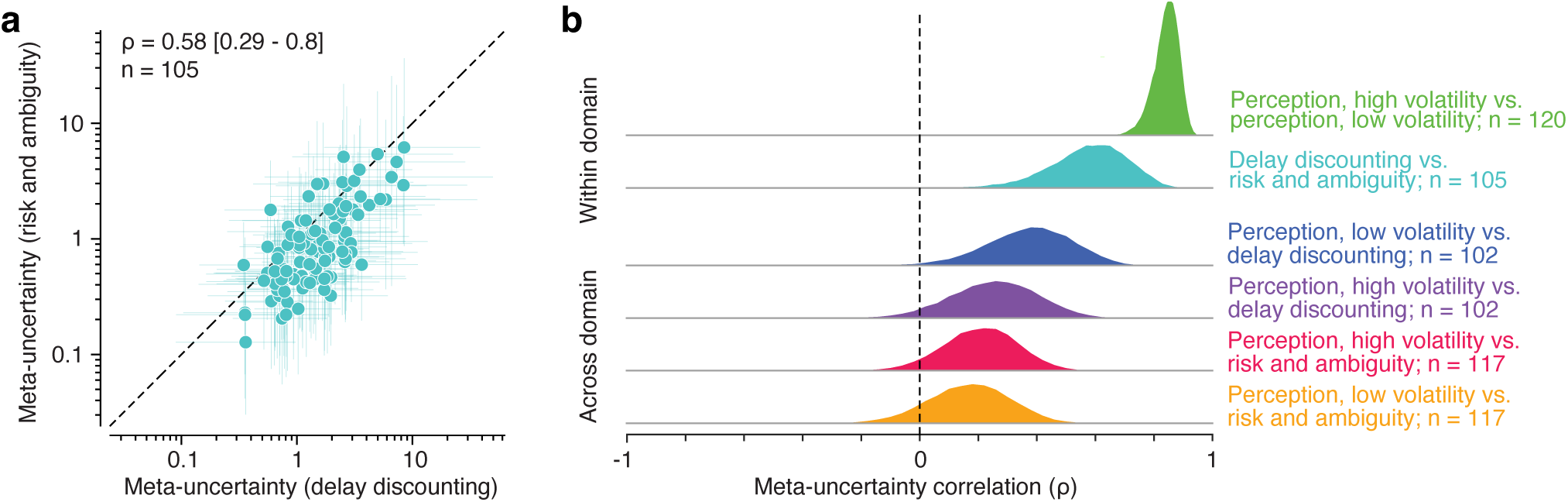
Metacognitive ability generalizes across tasks. **a**, Meta-uncertainty values estimated for the delay discounting task and risk and ambiguity task across 105 individual participants. Each symbol represents the geometric mean of the meta-uncertainty posterior for each participant in each task, and the vertical and horizontal lines represent the 95% credible interval in each dimension. The *ρ* value is that directly estimated from the model and not the result of correlating the point estimates. **b**, Posterior distributions of the correlation in meta-uncertainty across each task pair. The top two rows show decisive evidence for strongly positive within domain correlations in metacognitive ability. This is true for perceptual tasks (*ρ* = 0.84 [0.74 – 0.91]) and subjective decision-making tasks (*ρ* = 0.58 [0.29 – 0.8]). The bottom four rows show evidence for cross-domain correlations in metacognitive ability between perceptual and subjective decision-making. There is substantial evidence for a moderate correlation in the strongest of these comparisons and more equivocal evidence for the remaining three. From top to bottom: *ρ* = 0.37 [0.08 – 0.63], *ρ* = 0.25 [-0.05 – 0.53], *ρ* = 0.21 [-0.05 – 0.44], and *ρ* = 0.17 [-0.11 – 0.42]. The number of included

Our approach also confirmed previous reports of metacognitive generalization across perceptual tasks ^12,83–85^. When we fit the joint CASANDRE model to low and high volatility orientation discrimination blocks, we found a strong correlation in meta-uncertainty (Fig. 4b; *ρ* = 0.84, 95% CI = [0.74 *→* 0.91], Bayes Factor > 100, decisive evidence). The high correlation value likely reflects the similarity between the two tasks, which only differed in the number of randomly interleaved levels of contrast across trials (five for high and one for low volatility). We found little difference in the overall level of meta-uncertainty across the two tasks, in contrast to previous work suggesting that higher volatility in uncertainty across trials leads to higher meta-uncertainty ^46^.

Finally, we examined the domain generality of metacognitive ability by fitting the joint CASANDRE model to all combinations of tasks across perceptual and subjective decision-making. We found that the mean correlation value in meta-uncertainty across tasks was positive in all cases (Fig. 4b). However, of these comparisons only the meta-uncertainty correlation in the joint model of delay discounting and low volatility orientation discrimination reached a level of substantial evidence (Bayes Factor = 3.4). All other correlations did not achieve credible evidence (see Supplement). We conclude that our approach can effectively infer both the presence and absence of the conservation of metacognitive ability across tasks. While we found decisive evidence for conservation of metacognitive ability across subjective decisions, this ability does not necessarily generalize to highly distinct task domains or designs.

## Discussion

Our sense of confidence guides behavior to help us seek more information ^86^, refine our decision-making ^87^, and improve our outcomes ^88^. However, individuals vary in their metacognitive ability to channel and effectively use that sense of confidence ^5^. Current approaches to measuring metacognitive ability are limited in their conceptualization to tasks where confidence should relate to the probability of making a correct decision, and limited more practically in requiring many repeated trials of similar decision difficulty ^11,29,46^. Both of these limitations complicate efforts to assess an individual’s metacognitive ability in assigning confidence to their subjective decisions. Here, we overcome these hurdles by using a recently developed process model of confidence generation and extending it through Bayesian hierarchical fitting procedures ^46^. We find that the CASANDRE model provides a good account of metacognition in preference-based decision-making tasks, and further, that there is strong evidence that metacognitive ability generalizes across different kinds of subjective decisions.

Our work addresses an important and timely question in basic psychological and psychiatric research: whether individuals show consistent metacognitive abilities across different domains, behaviors, and contexts. This question is entangled with the more procedural one of how best to quantify and measure an individual’s introspective capabilities. While there are self-report instruments for this purpose ^89,90^, there has been growing interest in the development of task-based measures with good internal and external validity ^11,24,26,30,31,47,53,72^. The standard approach to operationalize metacognitive ability is to consider the accuracy with which individuals can discriminate correct from incorrect decisions with confidence reports ^11,29^. Task-based measures of metacognitive efficiency can minimize the influence of decision accuracy and criterion effects and have been used to great effect over the last decade. Many studies have established the potential external validity of these measures by finding relationships between the level of metacognitive efficiency and psychiatric conditions or symptoms where reduced introspection is a known feature ^8,33,91,92^. Likewise, many studies have established some degree of internal or construct validity of metacognitive efficiency by finding correlations across a variety of different tasks ^12,13,53^. An underlying assumption in this work is that confidence reports reflect an individual’s estimate of their probability of making a correct decision ^1^. As useful as that framework has been, it nonetheless has limited use for explaining confidence in internal mental processes where accuracy is not objectifiable ^24,46^.

Consider again the example of the car purchase decision. It is impossible to posit *ex ante* whether the correct (or best) choice is a high-end sports car or a station wagon for a given agent, unless you know what their (subjective) priorities are. When participants express their preferences through choices, we can model their decisions by inferring the utility they assign to the options ^93^. We can then further assess that a given choice may have violated the predictions of the utility model and that therefore the choice is “incorrect”. This notion of accuracy in preferential choice has allowed the metacognitive framework used in objective decision-making tasks to be applied to value-based decisions ^14,15,23^. However, this conceptually unsatisfying generalization of accuracy is likely part of the reason (along with technical limitations) why metacognitive ability has rarely been estimated from value-based decisionmaking tasks. A previous study by Folke et al. ^15^ calculated a measure of metacognitive accuracy in preferential decisions as the difference in the slope of psychometric functions conditioned on low or high confidence. A steeper slope indicates a stronger association between the participant’s choices and the difference in their valuation of the two options. Folke et al. found individual differences in this measure of metacognitive accuracy that correlated with other behaviors. However, this slope difference can be influenced by many factors other than metacognitive ability, including decision sensitivity and metacognitive bias, suggesting that fitting a computational model is required to isolate metacognitive ability ^46^.

Although most computational model-based measures of metacognitive ability rely on distinguishing correct from incorrect trials ^11,29,38^, there is a growing recognition that confidence reporting behavior often does not resemble an estimate of the probability of making a correct decision, even in objective decision-making tasks ^42,45,46^. When prior information or biases are introduced, participants will generally anchor their confidence reports to their choice consistency, instead of shifting reports to maximize the relationship between accuracy and confidence ^41–43,94^. Further, measuring the quality of confidence as the dissociation of correct and incorrect trials reduces confidence to error monitoring, which makes some sense for retrospective confidence judgments. However, both human and animal participants also exhibit highly lawful behavior when making a decision and confidence reports simultaneously ^31,39,67,95,96^. Instead, considering confidence as an estimate of noisy decision reliability provides a conceptually satisfying account of metacognitive ability that naturally applies as well to objective and subjective decisions. In this framework, participants estimate how likely they would be to make the same decision again ^15^, and metacognition is limited by the precision with which one can estimate uncertainty in an internal decision variable. In objective decision-making tasks, like perceptual discrimination, this uncertainty is expressed over the external variables to be discriminated. In value-based decision-making tasks, utility models provide a gateway into an individual’s subjective space, and the precision of their uncertainty estimate can be expressed over this internal subjective value difference. By combining utility modeling with the CASANDRE process model of confidence, we were able to account for confidence behavior across both task types and investigate the internal validity of metacognition for preferential decisions, where the stakes are real monetary rewards.

We find that metacognitive ability is highly conserved across different value-based decision-making tasks. Importantly, this relationship between meta-uncertainty estimated from a delay discounting task and a risk and ambiguity task did not appear to be driven by any similarity in decision-making behavior. There is a possibility that the noisiness of the internal representation of subjective value difference is related across both tasks ^82^, but we found little evidence that such a relationship could mediate the strong metacognitive generalization. Likewise, we found a very strong correlation between metacognitive ability estimated from two variants of an orientation discrimination task, confirming many previous reports of metacognitive generalization across perceptual tasks ^12,83,85,97^.

Similar to a growing body of literature, we find some evidence for domain generality in metacognitive ability. Previous work has found correlations in metacognitive ability estimated from tasks that vary widely, from perceptual discrimination in different sensory modalities (visual, auditory, somatosensory), to memory, and general knowledge ^12,13,53,98,99^. However, all these studies have focused on generality across tasks where there is an objectively correct answer, thus our finding across objective and subjective decisions is somewhat unprecedented. Further, many studies of domain generality make an effort to standardize all incidental aspects of task design to gain the best possible chance of finding a relationship in metacognitive ability ^13^. We took a different approach. If metacognitive ability is truly general and our estimation methods are accurate and powerful enough, we should observe correlations even using highly distinct task paradigms. Our perceptual tasks differed from our value-based tasks in nearly all aspects related to metacognitive measurement ^26,100,101^. In the value-based tasks, confidence was rated retrospectively on a four point scale with no incentives. In the perceptual tasks, a binary confidence report was made simultaneously with the decision, and reports were incentivized with different monetary rewards for high and low confidence decisions ^67^. Despite these substantial task differences, we find cross-domain correlations in meta-uncertainty near the level of cross-domain correlation in metacognitive efficiency reported in a large-scale, well-controlled study conducted by Lund et al. ^13^. Although the correlation estimates were comparable, their uncertainty was much greater in our results and only one out of four comparisons yielded substantial evidence for domain generality. This is likely because we collected data from less than a third of the participants and some of our tasks contained half the number of trials. This suggests that using our methods with a larger dataset would likely add to the credibility of the evidence for domain generality of metacognitive ability.

Our approach to investigating metacognitive domain generality relies on two interrelated innovations. The first is using value-based decision-making tasks where models for inferring the internal decision variable are well-validated, and the second is using a process model of confidence which can take into account and combine trials with different distributions of this decision variable. To illustrate why these considerations are critical, consider perception and memory, probably the most common domains where metacognitive generality has been tested ^12,13,53,98,99^. Although perceptual evidence can be well-controlled, on any given trial of a memory task, it is not really possible to know the strength of the internal evidence for recall. Though overall performance across memory trials may be controlled or matched to another task, this likely constitutes a mixture of individual trials that may be harder or easier. In contrast, each value-based decision-making trial is associated with an estimate of subjective value difference, and thus the signal strength used to make the choice. This property makes these economic decision-making tasks more similar to perceptual decision-making and more amenable to confidence modeling. The recently used numerical semantic memory task of Lund et al. ^13^ can be considered in the same way, and could in principle be employed without a staircase procedure if the resulting data were modeled using CASANDRE ^46^ or other similar process models of confidence generation ^47–50^.

We show that using Bayesian hierarchical modeling to explicitly model parameter correlations is critical for their accurate estimation ^13^. This is particularly true in experiments with relatively few trials and participants. This is one limitation of our study, and a potential reason for the limited evidence we find for domain generality. Our data collection was carried out online, and a portion of participants did not pass all quality checks. It will be important for future work to replicate our findings while potentially increasing the number of participants and trials per task, modifying task details, or performing a model comparison with other process models of confidence ^47,49,50^. On the other hand, the strong conservation in metacognitive ability within value-based decision-making tasks illustrates that these effects are robust to relatively small samples sizes. This is promising for the potential translation of this approach to clinical studies, where it is important to consider both the difficulty in recruiting and retaining patient samples, and the need to minimize the time burden placed on them.

We argue that subjective decisions are particularly well-suited to address many central questions in metacognition research. Numerous studies have documented alterations in metacognitive behavior or metacognitive ability in individuals suffering from mental health disorders ^8^. A real-world space where introspection about subjective processes often takes place is at the therapist or mental health clinician’s office. There, the fidelity of a client or patient’s report about their thoughts and behaviors is difficult or near-impossible to judge objectively. Metacognitive ability in this context could dictate not just how much veridical information the clinician has to work with, but also potentially determine the effectiveness of a given intervention. Value-based decisions such as risk-taking are central to many disorders in psychiatry ^58,102–105^, and assessing a person’s metacognition about this type of choice holds promise for the development of titrated interventions aimed at improving maladaptive attitudes toward risk. The CASANDRE framework lends itself well to implementation in a variety of other value-based tasks and decision models, such as reinforcement learning and other types of decision-making tasks that are relevant to psychiatry. Several recent papers have combined value-based decision-making tasks with metacognitive reports ^14,15,17,19^, and here we demonstrate a promising extension to this approach by developing a procedure for computationally evaluating metacognitive ability in subjective decision-making.

Building better models of decision quality is relevant for many applications, from psychiatry to artificial intelligence. Models that are systematically applicable to subjective choices — those that most resemble the highly consequential decisions we make every day — are even more relevant as we try to improve our understanding of how we introspect about our behavior, and how this process can go wrong. By combining value and confidence models, as we have done here, we indicate a path toward characterizing metacognition for many other types of decisions and behaviors that involve internally driven (subjective) processes, such as choices in interpersonal contexts, political choices, or financial decisions.

## Methods

### Study participants and Tasks

This study was approved by the National Institutes of Health Institutional Review Board. We recruited 128 participants (52 female) ages 18-55 (see Table 1 and Table 2 in Supplement for details on participant characteristics) from Amazon Mechanical Turk using CloudResearch’s MTurk Toolbox (formerly TurkPrime; see ^106,107^). Data collection took place between January and April 2023. Participants had to complete all three tasks to receive a participation payment. Choices were incentivized by providing a bonus in addition to the participation fee, corresponding to the outcome of one trial from either of the tasks, chosen at random at the end of the session (see Supplement). The tasks were programmed on PsychoPy (version 2022.2.4) using Pavlovia ((Open Science Tools, Nottingham, UK)).

In the perceptual decision-making task (Fig. 1c), participants were presented with a Gabor patch centered on the screen, with an image diameter set to 25% of the height of the monitor screen, and a spatial frequency of 6 cycles/image. On each trial the grating was rotated either clockwise or counterclockwise from vertical with one of eleven possible magnitudes of orientation angle between 0° and 5°. The task was composed of 4 blocks of 200 trials each. Two of the blocks corresponded to a low volatility condition with one stable level of contrast (0.06) across all trials, and the other two blocks corresponded to a high volatility condition with five different interleaved contrast levels (0.015, 0.03, 0.06, 0.12, 0.24). The task had a four-alternative forced choice design where participants had to report with a single button-press both the stimulus orientation (clockwise or counterclockwise from vertical) and their confidence level (high or low) (Fig. 1c). Participants were given feedback on their performance after their choice during the 5 initial practice trials but not in the rest of the task. Participants were told that one of the trials from the task (excluding practice) could be chosen at random at the end of the experiment and, if so, its reward outcome would be added to their payment as a bonus. The highest possible bonus corresponded to correct trials reported with high confidence and the lowest bonus was for incorrect trials reported with high confidence (see Supplement for details on payout structure). In each volatility condition, one block was a high incentive block and the other was a low incentive block that differed only in the potential payout of correct/high confidence trials (see “Task payment procedures” in Supplement for a detailed description of the payout matrix for each condition and for our bonus payment procedures). For the purposes of this manuscript, the high incentive and low incentive blocks within a volatility condition were pooled together. Block order was pseudorandomized as was orientation and contrast level within each block.

The delay discounting task (Fig. 1a) had 96 trials in a two-alternative forced choice design, where participants first made their choice between a smaller amount of money that they could receive that day or a larger amount of money that they could receive later. The immediate amount offered was $2, $10, or $20, the delayed amount ranged from $1 to $65. The delays ranged from 4 to 151 days. The amounts and delays changed from trial to trial, and the amounts were higher for the delayed offer in 90 of the trials, while in 6 trials ("catch trials") the immediate offer was larger than the delayed. The options were presented on alternating sides of the screen in a pseudorandom fashion. After making their choice, participants were asked to indicate their level of confidence that "they made the best decision for them" according to their preferences, on a 1-4 Likert scale. Participants were instructed that a confidence level 1 would mean they didn’t know if they chose what they preferred (e.g., chose randomly) while a confidence level 4 would mean they were certain they chose what they truly preferred.

Similarly, the risk and ambiguity task (Fig. 1b) had 80 trials in a two-alternative forced choice design, where participants first made their choice between a smaller amount of money that they could receive with a 100% probability or playing a lottery with a varying probability of winning a larger amount of money or $0^108^. 60 of the 80 trials were risk trials and the remaining 20 were ambiguity trials. On risk trials, probabilities varied from trial to trial and were 13, 25, 38, 50, or 75% and so did lottery amounts which ranged from $5 to $50. On ambiguity trials, the true lottery probability was 50% but 24, 50, or 74% of the probability information was masked. The expected value of the lottery was higher than that of the safe option in 60 of the trials, while in 18 catch trials it was strictly lower (first order stochastically-dominated) and in 2 catch trials it was equivalent (second order stochastically-dominated). The options were presented on alternating sides of the screen in a pseudorandom fashion. As in the delay discounting task, after making their choice, participants indicated their level of confidence that they chose what they truly preferred on a 1-4 Likert scale. Incentive compatibility was ensured by randomly selecting one trial and paying the participant according to their choice (see supplement).

To minimize the possibility of bots or disengaged participants in our dataset, the experiment was prematurely aborted if any of 3 conditions in any of the tasks were met: 1) More than 5% unanswered trials, 2) identical single button-presses for all trials, and/or 3) poor performance on catch trials (see Supplement for details). Once the experiment was complete and prior to running task comparison analyses, participant datasets were evaluated for data quality (see Supplement).

### Utility modeling of subjective decision task behavior

All models were fit using Stan (command line interface version cmdstan 2.30.1) (https://mc-stan.org/). The code package for the utility and CASANDRE models can be found in our Github repository (https://github.com/CDN-Lab/CAStanDRE).

We fit two utility models to the trial-by-trial choice behavior in the subjective decision-making tasks. For the risk and ambiguity task, we fit the following model ^74^:

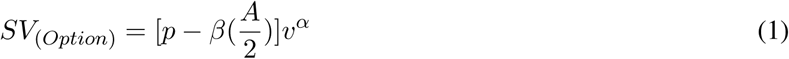

where *p* is the probability of the option, *A* is the ambiguity level (proportion of the probability information that is hidden) and *v* is the amount (in US $). *α* and *β* are the free parameters of the model and represent a subject’s risk tolerance and ambiguity tolerance, respectively. *β <* 1 corresponds to risk aversion and *β >* 1 corresponds to risk seeking preferences. *α >* 0 corresponds to ambiguity aversion and *α <* 0 corresponds to ambiguity seeking preferences.

For the delay discounting task, we fit the following model of discounted utility ^65^:

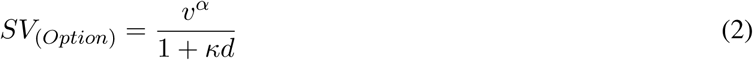

where *v* is the amount (in US $), *d* is the delay (in days). *α* is the risk tolerance obtained by fitting model 2 to data from the subject’s choices in the risk and ambiguity task, while *κ* a free parameter corresponding to the discounting rate. Higher *κ* values indicate a higher rate of reward devaluation as a function of time.

Subjective values of the options in the subjective decision-making tasks were compared and converted to choice according to the following decision rule ^75,76^:

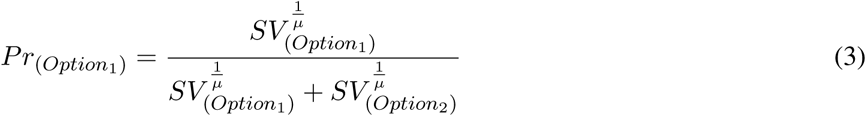

where the free parameter *µ* (*µ_risk_*for risk and ambiguity task and *µ_delay_*for delay discounting task) represents choice stochasticity, with higher values indicating lower sensitivity of the decision to the subjective value comparison.

We hierarchically fit participant-level as well as group-level parameters. Strictly positive parameters (*α*, *κ*, *µ_risk_*, and *µ_delay_*) were modeled using hyperparameterized lognormal distribution priors, while *β* was modeled using hyperparameterized normal distribution priors. We employed empirical data ^65^ to center the distributions of the *α* (0.65) and *κ* (0.05) hyperpriors. *β* was the only parameter with empirically determined bounds, between *±*1.35, constraining the modeled probability in the risky and ambiguous utility model, 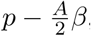, to be between 0 and 1.

### CASANDRE model

We model confidence in choice behavior using the confidence as a noisy decision reliability estimate (CASANDRE) model ^46^, in which the confidence variable, *V_c_*, is modeled as:

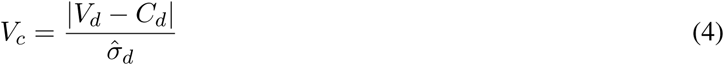

That is, a signal to noise ratio where signal strength is computed as the distance between the decision variable *V_d_* and the decision criterion *C_d_*, normalized by the subject’s estimate of their decision uncertainty, 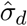. This is compared to an internal confidence criterion, *C_c_*, a free parameter of the model, to obtain the confidence report. Confidence criteria were allowed to be asymmetrical for all tasks, that is, different for each type of choice (e.g., left versus right in the perceptual task, or lottery versus safe in the risk and ambiguity task). This generally allowed for better fits, especially in the value-based tasks ^46^.

The decision is based on the value of a decision variable, *x*, that can be considered in units of an external stimulus feature for perceptual decisions or as the difference in subjective value between two options for a subjective decision. CASANDRE assumes that a participant is on average unbiased, but does not have direct access to *x*. Instead they can estimate a noisy representation of it on a trial-by-trial basis that depends on their decision sensitivity, modeled by the inverse of the free parameter, *ω_d_*.

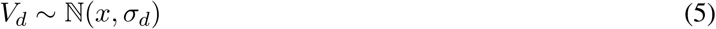

The lower this free parameter, the higher the decision sensitivity, and the more accurate the internal representation of *x* is on each trial. CASANDRE likewise assumes that a subject does not have direct access to the level of noisiness of their internal representation, *σ_d_*, but only an estimate of it, 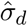.

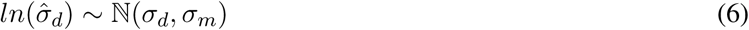

This estimate is strictly positive and modeled as a trial-by-trial sample from a lognormal distribution whose mean is determined by the true noisiness on the decision variable, with spread depending on a subject’s uncertainty about this noise, modeled with the free parameter *σ_m_*, termed meta-uncertainty, where higher values indicate lower metacognitive ability.

### Joint hierarchical Bayesian Copula models of CASANDRE and meta-uncertainty correlations

We developed a Bayesian hierarchical extension to the CASANDRE likelihood model in Stan to simultaneously fit the CASANDRE parameters and the meta-uncertainty correlations across tasks. For each task comparison, we fit participant-level as well as group-level parameters, and a single Copula-based correlation parameter. This correlation parameter was used to probe the relationship between the participant-level meta-uncertainty parameters, CASANDRE’s measure of metacognitive ability, across tasks. Given the computational complexity of this model, it was optimized for multithreading and was run on our high performance cluster, Biowulf.

For the perceptual decision task, five parameters of interest for each participant were sampled hierarchically: decision sensitivity, decision criterion, meta-uncertainty, and confidence criterion for the "clockwise" and "counter clockwise" options. Additionally, each participant’s lapse rate was sampled non-hierarchically. In the high volatility blocks, there were five contrast levels, each fitted with their own decision sensitivity and decision criterion; and two confidence contexts (low vs high incentive), each fitted with their own "clockwise" and "counter clockwise" confidence criterion.

For the subjective decision tasks, the CASANDRE parameters were similarly sampled hierarchically but without the decision criterion, which was constrained by the prior fitting of the utility models to be 0. Asymmetric confidence criteria were sampled in these tasks as well, but corresponded to the "now" and "later" options in the delay discounting task, and the "safe" and "risky" options in risky and ambiguous task.

Every hierarchically sampled parameter of interest had group level *µ* and *ω* hyperparameters, in each of the two tasks being compared. Strictly positive parameters in CASANDRE models were modeled using hyperparameterized lognormal distribution priors, while all other parameters, with the exception of lapse rate in the CASANDRE model and the *ρ* correlation parameter, were modeled using hyperparameterized normal distribution priors. Lapse rate was modeled using a beta distribution prior with a mean and mode of 0.015, reflecting the small number of expected errors in these experiments. *ρ* was modeled with a normalized uniform distribution prior with bounds -1 and 1. For our hyperpriors, we selected standard normal distributions for the location parameters of the priors, and standard lognormal distributions for the scale parameters of the priors.

To model the dependency structure between meta-uncertainty samples across each pair of tasks, we used a Copula model. One Copula-based correlation parameter was sampled per comparison. This model takes the meta-uncertainty parameter samples from each task, as well as the sampled free parameter, *ρ*, as inputs. The meta-uncertainty parameter samples are first converted into their cumulative densities through a lognormal CDF (with the location and shape hyperparameters from each task). These cumulative densities are transformed into a standard normal distribution through the inverse standard normal CDF. Finally, these standard normal distributions are compared using a bivariate normal distribution PDF with a correlation coefficient equal to the sampled correlation parameter, *ρ*.

### Parameter recovery

Parameter recovery was carried out for the perceptual and value-based models independently, matching previously estimated parameter range and number of observations: *N_within perceptual_* = 120, *N_within value_* = 105, *N_perceptual vs risk_* = 117, *N_peceptual vs delay_* = 102. In MATLAB (version R2023b), generative parameters were sampled at random from bounded prior-matched distributions for all participant-level CASANDRE model parameters. For parameter recovery of the subjective decision tasks, decision criterion was excluded from the generative parameters. Generative parameters were then used to create synthetic data sets from the CASANDRE likelihood function. Task structures and trial numbers were matched in the synthetic data sets to each of the experimental tasks, with perceptual task data utilizing separate decision sensitivity and decision criterion for each of the five contrast level, and separate asymmetric confidence criteria for each of the two confidence contexts.

Synthetic data was then packaged and run on each model independently using Stan to generate posterior distributions over all parameters. The log of the geometric means of the strictly positive posterior distributions were compared in scatter plots to the log of the ground truth generative parameters, checked for convergence to the line of unity, and inspected for bias by the slope of their Deming regression lines. For all other parameters, the means of the posterior distributions were compared in scatter plots to the ground truth generative parameters, again for convergence to the line of unity as well as through inspection of the slope of the Deming regression line. All parameters of interest of all models were recovered without systematic biases.

### Correlation recovery

To evaluate the correlation of meta-uncertainty parameters across tasks we compared two approaches. The first method (“separate”) was to fit meta-uncertainty for each participant in independent hierarchical models for each task, calculate point estimates from the resulting posterior distributions, and then correlate this set of point estimates using Spearman’s *ρ*. The second was to directly build a dependency structure, in our case a Copula model, into the sampling program model, which we estimated jointly with the CASANDRE parameters (see Section Joint hierarchical Bayesian Copula models of CASANDRE and meta-uncertainty correlations).

To identify which method provided the best estimate of the correlation in metacognitive ability across tasks, we performed a correlation recovery analysis for both approaches. For the generative model, we drew meta-uncertainty values from a multivariate lognormal distribution matching the model prior. We simulated 21 sets of meta-uncertainty values per task, with equally spaced underlying correlation coefficients, drawn from this multivariate distribution for each synthetic participant in each confidence condition for the two tasks being compared. This returned roughly equally spaced ground truth correlation coefficients between approximately 0 and 1. We matched the number of participants and trials to that of the tasks to be compared.

The compute the ground truth correlation coefficient we fit a lognormal distribution to each set of generated meta-uncertainty parameters to obtain the distribution’s *µ* and *ω* parameters. We then calculated the cumulative densities of each of the meta-uncertainty parameters using a lognormal CDF with these fitted *µ* and *ω*. We obtained the normalized value of these cumulative densities by calculating their inverse standard normal CDF values. Finally, we computed the Pearson’s r between the two transformed distributions. This mirrors how the Copula in the joint models work.

The synthetic data generated by these and the other independently drawn model parameters were then fit separately with the “separate” and “joint” models. For the “separate” model, we used the geometric mean of each independent model’s posterior distribution as the parameter’s point estimate to compute the recovered correlation coefficient. For the “joint” model, we used the mean of the posterior distribution for *ρ* as the recovered correlation coefficient.

## Acknowledgements

This research was supported (in part) by the Intramural Research Program of the National Institute of Mental Health (ZIA-MH002988, PI SL-G) and the NEI, and by U.S. National Institutes of Health grant K99 EY032102 (to CMZ).

We thank Cassie Raymond, Savana Agyemang, and Santiago Guardo-Maya for help in conducting the online experiments, and Ricardo Pizzaro for help in developing analysis software. We thank Bruno Averbeck for helpful conversations and support on this project. This work utilized the computational resources of the NIH HPC Biowulf cluster (https://hpc.nih.gov). The development of the models used in this paper was made possible in part by the generous guidance of Stan’s developers and other users on the Stan website’s modeling forum (https://discourse.mc-stan.org.

## Author contributions

CMZ and SL-G conceived the study. CMZ, SL-G, and ZMB-S designed the study. DG collected data. CRP, CYZ, CMZ, and SL-G analyzed data. CRP, CMZ, and SL-G wrote the first draft. All authors edited the paper.

## Competing interests

The authors declare no competing interests.

## Notes

https://github.com/CDN-Lab/CAStanDRE

### Competing Interest Statement

The authors have declared no competing interest.

