## Supplement for "Computational characterization of metacognitive ability in subjective decision-making"

Supplementary material

### Participant characteristics

**Table 1** **Demographic characteristics of the study sample**

| | Number | % |
|---|---|---|
| **Age (Years)** | | |
| 18 - 23 | 4 | 3.1 |
| 24 - 29 | 17 | 13.3 |
| 30 - 35 | 33 | 25.8 |
| 36 - 41 | 35 | 27.3 |
| 42 - 47 | 16 | 12.5 |
| 48 - 55 | 23 | 18.0 |
| **Race and Ethnicity** | | |
| Asian | 12 | 9.4 |
| Black, African American | 18 | 14.1 |
| White | 91 | 71.1 |
| Multiracial | 7 | 5.5 |
| Hispanic | 12 | 9.4 |
| **Education Level** | | |
| None completed | 1 | 0.8 |
| Diploma or GED | 38 | 29.7 |
| Associate or Trade Diploma | 18 | 14.1 |
| Bachelors | 59 | 46.1 |
| Masters | 10 | 7.8 |
| Doctorate | 2 | 1.6 |
| **Income (Annual)** | | |
| $0 - $9,999 | 10 | 7.8 |
| $10,000 - $15,000 | 9 | 7.0 |
| $15,000 - $24,999 | 18 | 14.1 |
| $25,000 - $49,999 | 36 | 28.1 |
| $50,000 - $99,999 | 42 | 32.8 |
| > $100,000 | 13 | 10.2 |

Values are frequencies and percentages.

**Table 2** **Mental health characteristics of the study sample**

| | Number | % |
|---|---|---|
| **History of Anxiety or Depression** | | |
| Yes | 39 | 30.46 |
| **History of Substance Use** | | |
| No smoking | 61 | 47.65 |
| Fewer than 5 cigarettes per day | 27 | 21.09 |
| 6-10 cigarettes per day | 13 | 10.15 |
| 11-20 cigarettes per day | 23 | 17.96 |
| >20 cigarettes per day | 4 | 3.12 |
| No alcohol | 62 | 48.43 |
| Fewer than 5 drinks per month | 35 | 27.34 |
| 6-10 drinks per month | 14 | 10.93 |
| 11-20 drinks per month | 7 | 5.46 |
| >20 drinks per month | 10 | 7.81 |
| History of treatment for substance use | 7 | 35.46 |

Values are frequencies and percentages.

### Task payment procedures

All participants who were offered $20 as a participation payment to be disbursed via the study platform payment at the end of the study. In addition, we incentivized choice behavior in the tasks by offering an additional bonus as is described below. Participants were told that both participation payment and bonus would be contingent on the completion of all assigned tasks and questionnaires.

Specifically, subjects were instructed that at the end of the study one of the choices they made on one of the tasks would be randomly chosen and their bonus reward will correspond to the outcome of that choice. For example: 1) if the random trial selected corresponded to the perceptual decision task, the bonus would depend on the type of block (high incentive or low incentive) and whether the answer was correct or incorrect and the confidence reported was high or low according to the following payout matrix (table 3):

| | Low Incentive Blocks | | High Incentive Blocks | |
|---|---|---|---|---|
| | Correct | Incorrect | Correct | Incorrect |
| High Confidence | $9 | $0 | $13 | $0 |
| Low Confidence | $8 | $6 | $8 | $6 |

**Table 3** perceptual task incentivization structure by incentive blocks.

2) if the random trial selected corresponded to the risk and ambiguity task, the bonus would depend on whether the participant chose the "safe" option or the "lottery" option. If the "safe" option was chosen the bonus would be $5. If the "lottery" was chosen, then the lottery would be automatically actualized. This meant that the computer would randomly draw a color (red or blue) from the true distribution corresponding to that trial. The resulting color would

determine whether the outcome was a monetary gain or \$0. The bonus would correspond to that value. 3) if the random trial selected corresponded to the delay discounting task, the bonus would depend on whether the participant chose the "smaller sooner" or the "larger later" reward. For example, the task could be choosing between \$5 now or \$65 in 29 days. If the "smaller sooner" was chosen, the bonus would correspond to \$5 and would be made available to the participant via the Cloudresearch payment the same day they completed the task. If instead the "larger later" was chosen, the bonus in this example would correspond to \$65 and would be made available via CloudResearch payment on the date that corresponded to the number of days from the date that the task was completed.

This compensation mechanism was made clear to participants with thorough instructions. Critically, there was no deception as to the mechanism of determining each participant's bonus.

### Premature termination of experiment and behavioral data quality control

Prior to task comparison analyses, we evaluated each participant dataset as to whether it exhibited enough confidence variability in each task that would be compared, in order to adequately fit the CASANDRE Copula model. We excluded datasets from a comparison if the participant answered with the same confidence level on at least 95% of trials in any of the tasks. The number of exclusions differed across comparisons (8 for low volatility vs high volatility; 23 for delay discounting vs risky and ambiguous decison-making; 26 for perceptual vs delay discounting; and 11 for perceptual vs risky and ambiguous decision-making).

### Delay discounting and risky decision making behavior

To minimize the possibility of bots in our dataset, the experiment was prematurely aborted if any of 3 conditions in any of the tasks were met: 1) More than 5% unanswered trials, 2) identical single button-presses for all trials, and/or 3) poor performance on catch trials. For the perceptual task, the catch trials corresponded to the trials with the highest contrast and strongest orientation for each block. In the low volatility blocks, the stopping criterion was met when the response was accurate in fewer than 14 of the 20 catch trials. In the high volatility blocks, the stopping criterion was met when the response was accurate in fewer than 6 of the 8 catch trials. The probability of responding randomly and passing these criteria was lower than 15%. For the risk and ambiguity task, the stopping criterion was met when the response was the stochastically dominant option in fewer than 10 of the 20 catch trials (18 had first-order and 2 had second-order stochastically dominant options). The probability of responding randomly and passing this criterion was lower than 1%. For the delay discounting task, the stopping criterion was met when the response was the delayed option in 5 of the 6 trials where the delayed option amount was less than the immediate reward. The probability of responding randomly and passing this criterion was lower than 4%.

The utility parameters showed adequate diversity in our sample (see Fig.S1). The mean discount rate ($\kappa$) was 0.0084 ($ln(\kappa) = -4.8$) which means that on average monetary rewards lost only approximately 23.5% of their value in 365 days (1 a). This annual discount rate is comparable to but lower than what has been typically described in the literature (33% according to[1]). In parallel, participants exhibited a slightly higher level of risk aversion ($\alpha = 0.56$) (Fig.S1 b) compared to similar studies (see[65]).

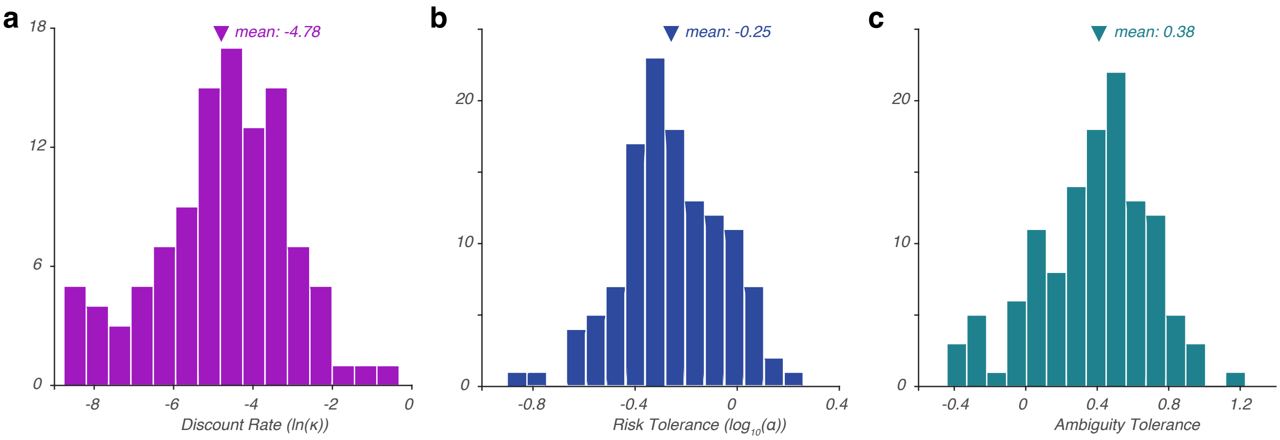


**Supplementary Figure 1** Utility model parameter distributions

We found no correlation between the general preference for lotteries and for immediate rewards (correlation between the % of lottery choices and the % of immediate options was not significant, $\rho = 0.16$, $p = 0.1$). However, the discount rate was significantly correlated with the risk tolerance parameter such that more impatient individuals were also more risk seeking ($\rho = 0.43$, $p < 0.001$). Note that this is not explained by confounds in the estimation of the discount rate due to non-linear utility ([3]) as discount rates were estimated after taking risk aversion levels into account using a non-linear utility discounting model ([65]). Risk tolerance ($\alpha$) and ambiguity tolerance ($\beta$) were uncorrelated ($\rho = 0.11$, $p = 0.22$), as were the discount rate and ambiguity tolerance ($\rho = 0.02$, $p = 0.83$). Overall, the differentiation and diversity of these utility parameters supports the idea that these are dissociable decision-making constructs, as is well established in the literature.

### Meta-uncertainty and decision sensitivity parameters

Similarly, we found that the CASANDRE parameters, decision sensitivity and confidence sensitivity exhibited significant diversity in this sample (Fig. S2). Decision sensitivity was generally higher for the risky decision-making task (mean $1/\sigma_d = 1.4$) compared to the delay discounting task (mean $1/\sigma_d = 1.2$) and the perceptual decision-making task (mean $1/\sigma_d = 0.8$). Conversely, meta-uncertainty (inverse of confidence sensitivity) was generally worse in the subjective decision-making tasks (mean $\sigma_m = 1.9$ for delay discounting and mean $\sigma_m = 1.1$ for risky decision-making) than in the perceptual decision-making task (mean $\sigma_d = 0.5$).

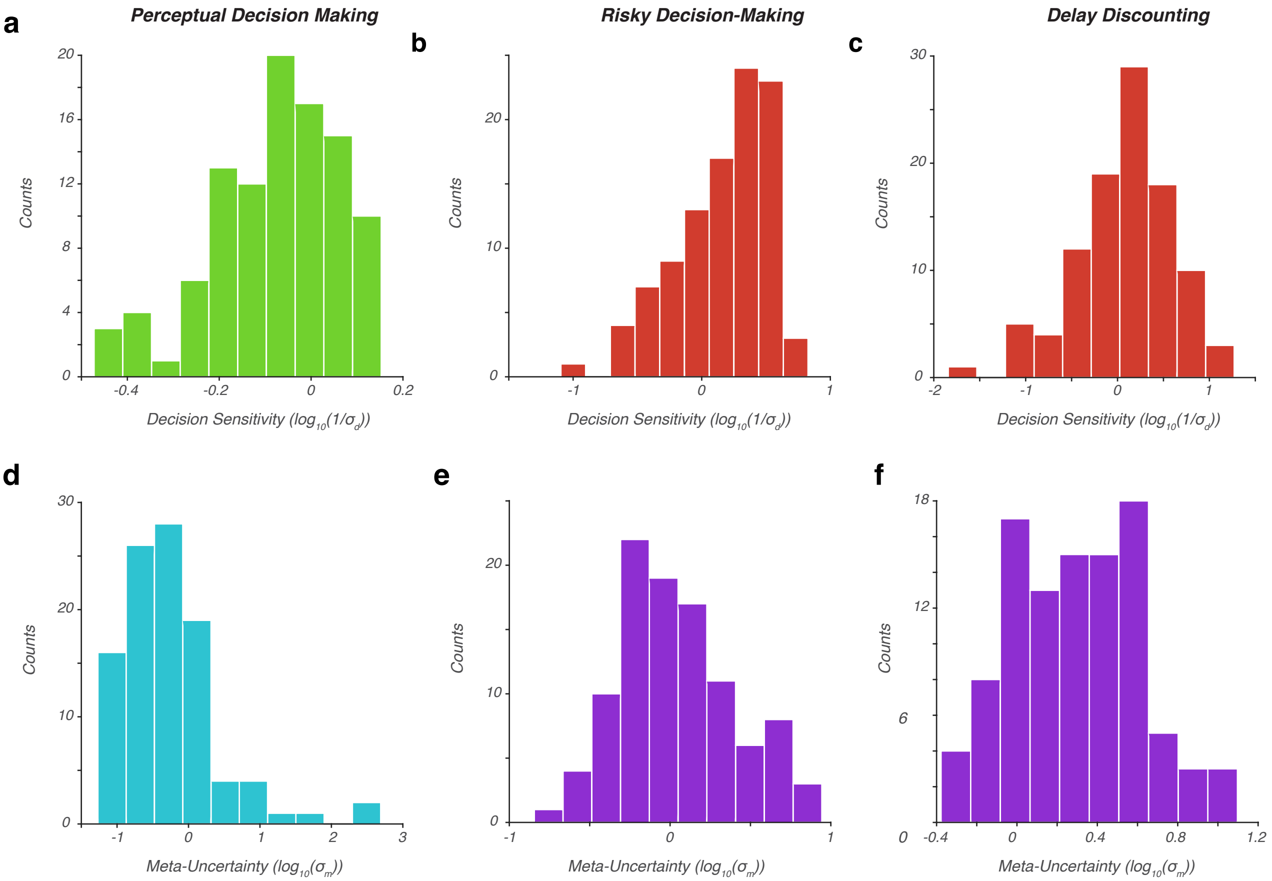


**Supplementary Figure 2** CASANDRE parameter distributions (Meta-uncertainty and decision sensitivity)

### H-Meta d' model M Ratio estimates

We used a Bayesian hierarchical modeling approach to estimate the correlation in metacognitive ability between the subjective decision-making tasks utilizing an alternative measure of metacognitive sensitivity, the M-Ratio[29]. For this analysis we used the H-Meta d' code package (`https://github.com/metacoglab/HMeta-d`) to fit a descriptive signal detection theory model of metacognition[71]. As in our main analysis, we estimated a correlation coefficient ($\rho$), to our delay discounting and risky and ambiguous decision-making task data sets. Compared with the copula modeling approach, this estimates the correlation of metacognitive ability across tasks by drawing log transformed M-Ratio samples for each task from the same bivariate normal prior, where the correlation coefficient ($\rho$) of the prior is sampled as a hyperparameter from a uniform hyperprior. The same exclusion criteria were applied for this comparison as in our model, with participants showing less than 5% variability in their confidence choices dropped from the comparison (number removed = 23). In order to apply the meta d' model, we coded all trials with the same sign of subjective value difference together as one stimulus level. For example, all trials were the utility model predicted that the safe option would have a higher subjective value than the lottery were pooled together, and vice versa.

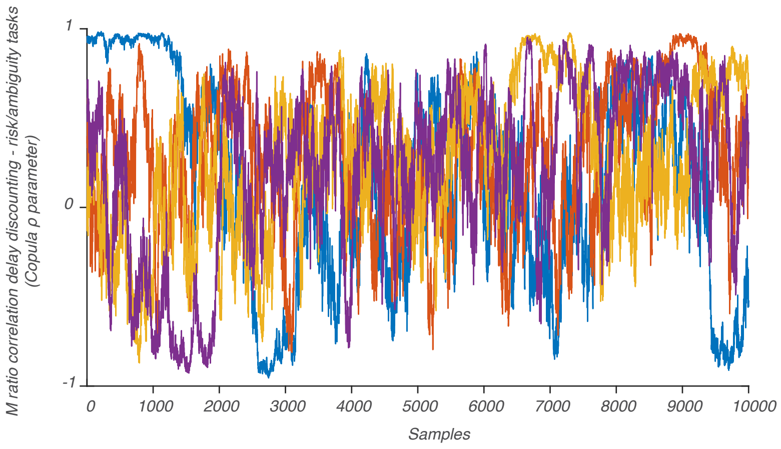


**Supplementary Figure 3** H-meta d' M-ratio model

We noted that the correlation parameter failed to visually converge (see trace plot Fig.S3), with parameter samples that spanned from nearly perfect anti-correlation to nearly perfect correlation. We speculate that this is because the underlying subjective value difference distributions of the value-based tasks are reduced to a binary for the purposes of this model (as in "stimulus 1" and "stimulus 2"). We conclude that these tasks are therefore not suitable for fitting by naively applying a descriptive model measure like M-Ratio.

### Additional model fitting details

The number of free parameters depended on the number of subjects included in each task comparison Copula model, as well as the number of contrast levels and confidence contexts:

| Comparison | Number of observations | Number of parameters |
|---|---|---|
| low volatility vs high volatility | 120 | 2561 |
| delay discounting vs risk and ambiguity | 105 | 1059 |
| low volatility vs delay discounting | 102 | 1349 |
| high volatility vs delay discounting | 102 | 2181 |
| low volatility vs risk and ambiguity | 117 | 1544 |
| high volatility vs risk and ambiguity | 117 | 2496 |

**Table 4** number of participants included, and free parameters fit, in each task comparison Copula model.

### Model credibility analysis

Bayes factor quantifies the amount of evidence for or against an alternative hypothesis when compared to the null hypothesis. When one is considering $BF_{10}$, the ratio of the posterior probability of $H_1$ over $H_0$, 1 represents equivocal evidence while values much greater than 1 indicate strong evidence for the alternative hypothesis. Calculating this ratio can be tricky, but, fortunately, there is a simple method that works even for hierarchical models, the

Savage-Dickey density ratio[6]. It is given by the formula:

$$BF_{10} = \frac{p(D|H_1)}{p(D|H_0)} = \frac{p(\phi = \phi_0|H_1)}{p(\phi = \phi_0|D, H_1)} \tag{7}$$

In essence, this is a comparison of models — the model specified by the data, given the likelihood function and the prior, versus the null model specified only by the prior. In the case of correlations, this is the probability density function of the prior at 0, $p(\phi = \phi_0|H_1)$, and the probability density function of the posterior at 0, $p(\phi = \phi_0|D, H_1)$. Fig.S4 shows the Bayes factor calculated using the Savage-Dickey density ratio for meta-uncertainty correlation in each task comparison. For both within-perceptual ($BF_{10}$ > 1000) and within-value ($BF_{10}$ = 157) the evidence for a correlation is decisive, while the evidence for a correlation between the low volatility blocks of the perceptual task and the delay discounting task ($BF_{10}$ = 3.4) is substantial, per the Kass and Raftery table[7]. The Bayes factors for the remaining comparisons trend in the opposite direction but do not rise to the level of substantial evidence for the null hypothesis.

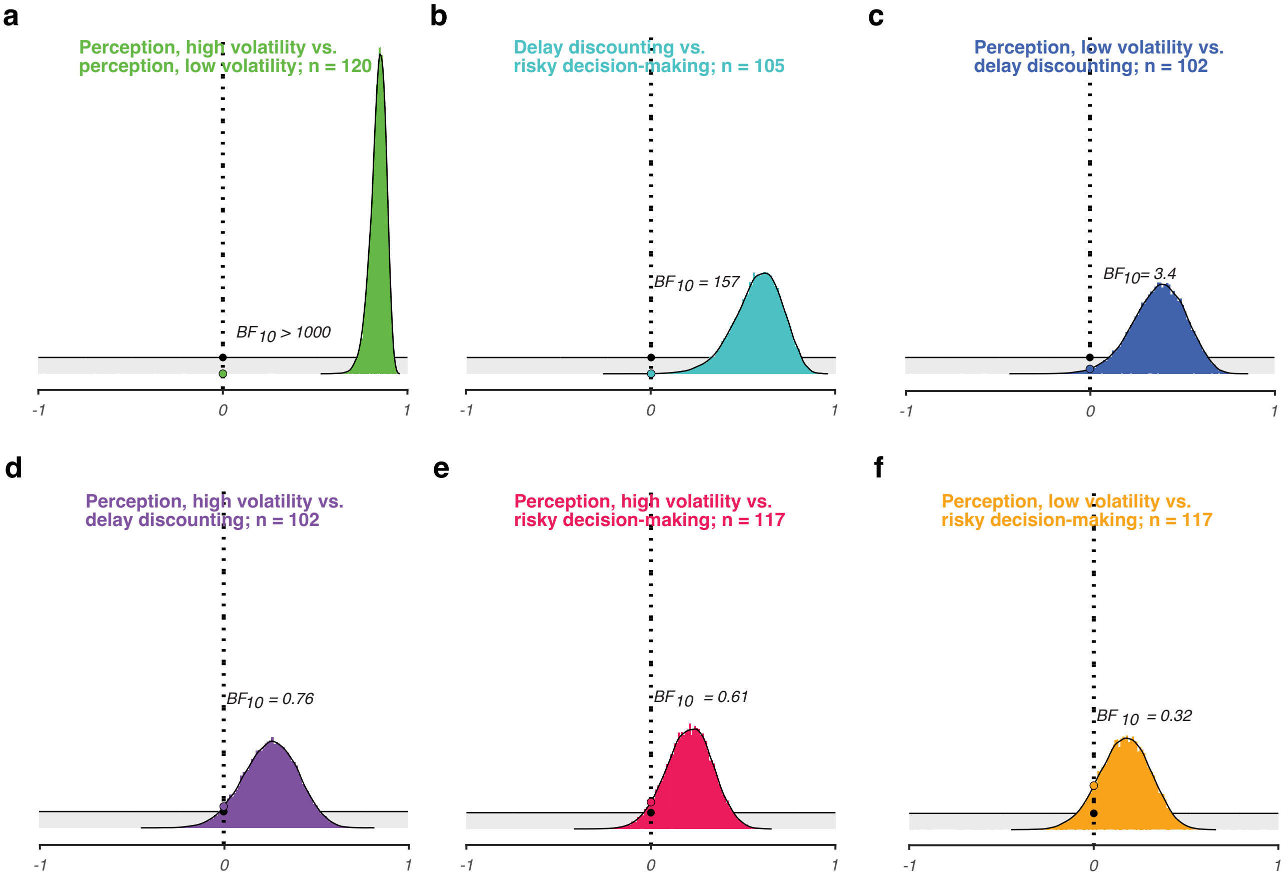


**Supplementary Figure 4** Savage-Dickey Density Ratio.

Figure S4. Bayes Factor Analysis for modeled correlations

### Posterior checks

Conclusions drawn from posterior distribution samples are only trustworthy when certain conditions are met. Sampling should be comprehensive, converge to a typical set, minimize sampling error, and show good parameter mixing. We employed a mixture of statistical and visual checks to ensure that this was the case for all of our posteriors at the participant, group, and correlation levels.

Our quality checks included: 1) verifying adequate sampling by inspecting effective sample size, 2) confirming statistical convergence through Gelman-Rubin's $\hat{R}$ measures, 3) inspecting that each chain of our posteriors was well explored by comparing the marginal energy distribution to the transitional energy distribution, 4) ruling out sticking behavior or skewed rank distributions by observing rank plots, and 5) confirming good convergence and mixing in trace plots.

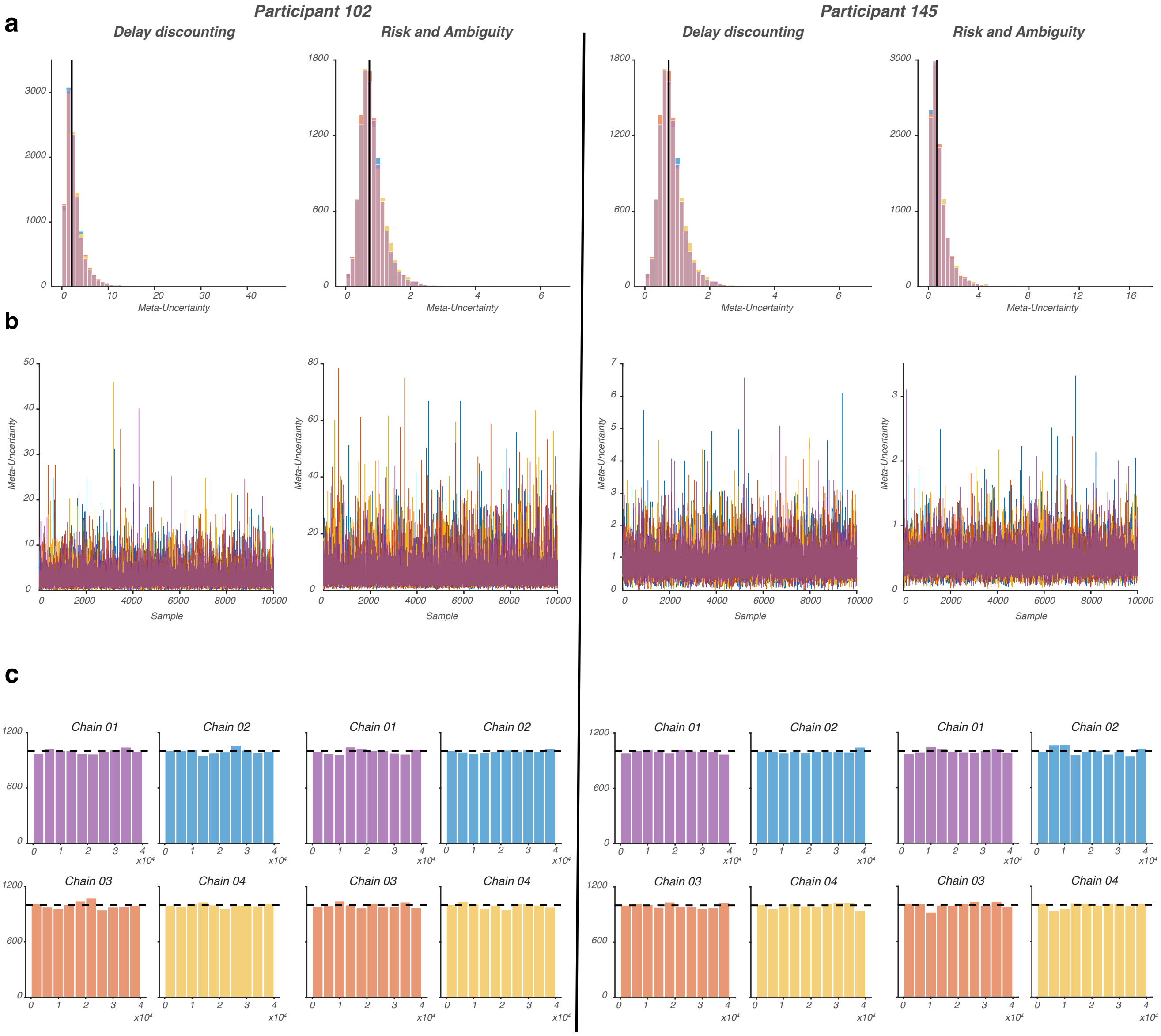


**Supplementary Figure 5** Example participant diagnostic plots

Figure S.5 depicts some diagnostics corresponding to the meta-uncertainty parameter estimation in the example participants shown in Fig. 2. Parameter histograms (Fig. S5 a) show significant overlap of the 4 chains and good estimation by geometric mean, and trace plots (Fig. S5 b), generated by creating line plots showing the change in parameter value on each draw, showed adequate convergence and stability, with a few excursions of parameter values but no systematic issues. Rank plots (Fig. S5 c) made by combining the chains, ranking their values, and then separating the ranks back into their original chains, are the most sensitive test of convergence. Rank plots showed uniform distributions within each chain, which signals good parameter mixing, convergence, and good posterior exploration.

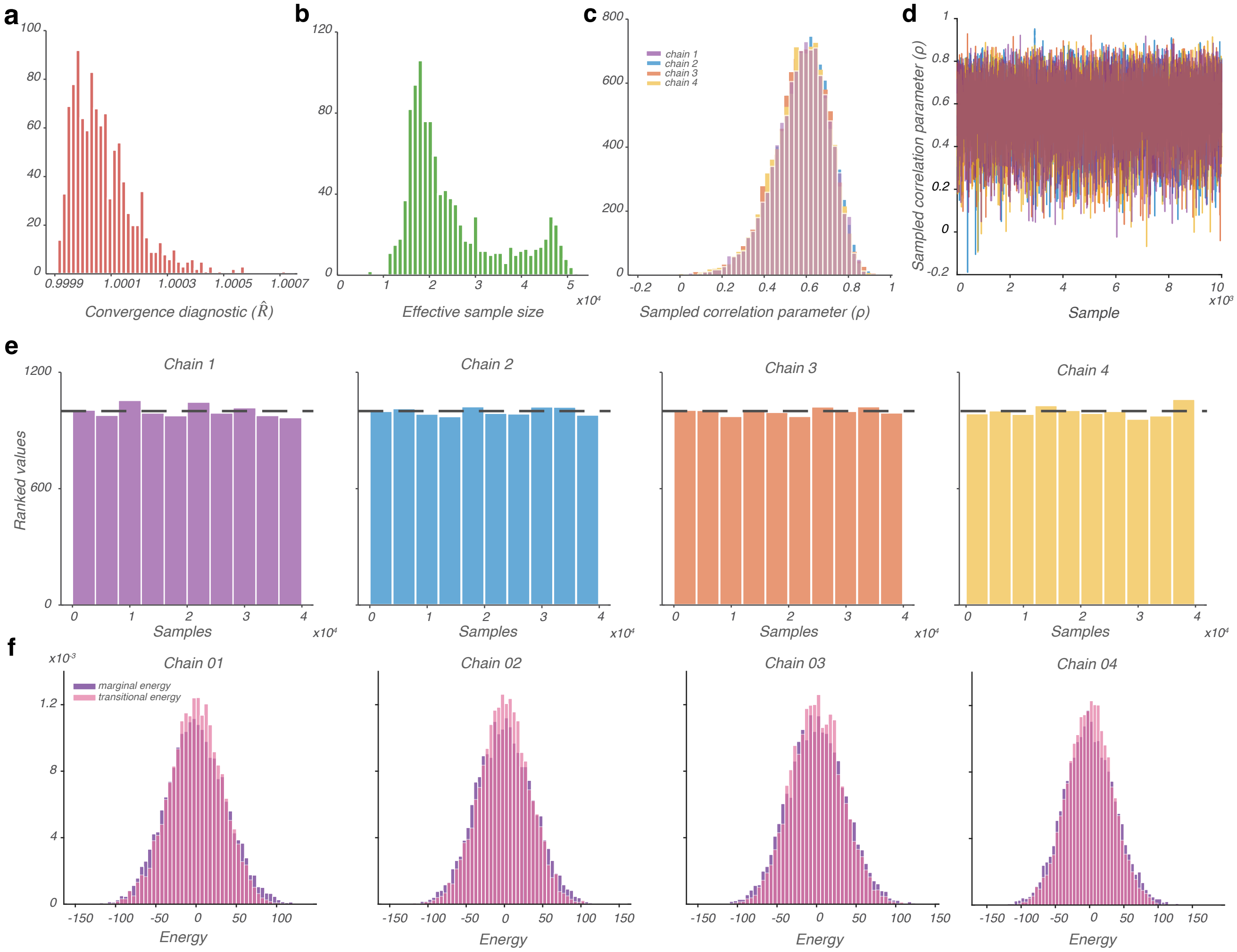


**Supplementary Figure 6** Diagnostic plots for copula model of correlation between delay discounting and risk and ambiguity meta-uncertainty

Figure S.6 depicts all diagnostics for the main analysis of the delay discounting versus risk and ambiguity meta-uncertainty correlation. The parameter of interest here is the copula correlation coefficient ($\rho$). Gelman-Rubin's $\hat{R}$ (Fig. S6 a) was between 0.99 and 1.01 for all model parameters, which is the criteria for statistical convergence. Effective sample size (Fig. S6 b) was greater than 400 for all model parameters, as would be expected for efficient and comprehensive sampling. The $\rho$ parameter distribution (Fig. S6 c) overlapped greatly across all four chains, and the trace plot (Fig. S6 d) confirmed good convergence and very good stability of the parameter values. Rank plots (Fig. S6 e) showed uniform distributions within chains. Finally, we compared the marginal energy distribution to the transitional energy distribution (Fig. S6 f). The marginal energy distribution is the mean corrected distribution of the energy of the sampler on each draw, while the transitional energy represents the change in energy on each draw. These two distributions were found to have nearly perfect overlap, signaling full exploration of the posterior in the sampling process.

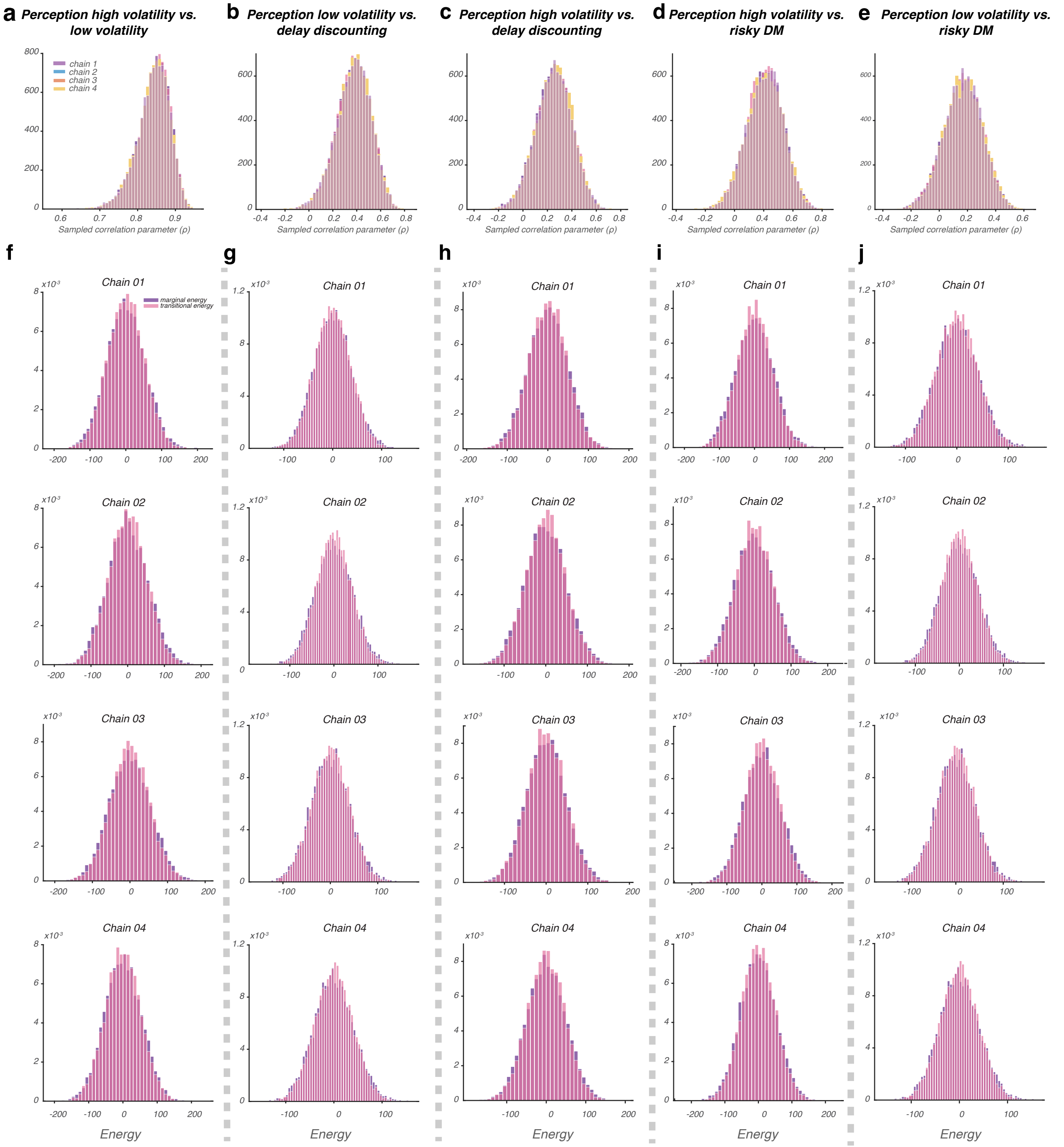


**Supplementary Figure 7** Diagnostic plots for copula model of correlation between all other tasks

Model fitting quality was similarly verified for all other copula models. $\rho$ parameter distributions for the other within-domain comparison (perceptual decision-making task high volatility versus low volatility blocks) (Fig. S7 a), and the cross-domain comparisons (Fig. S7 b through e) showed overlap across chains and high stability. Similarly, full exploration of the parameter space during sampling was verified by energy plots for each chain.

#### Predicting meta-uncertainty from decision sensitivity with generalized linear models

We ran generalized linear mixed-effects models with random intercepts and slopes per participant and tested the relationship between meta-uncertainty parameters and decision sensitivity parameters across tasks (see Methods). Model 1 predicted meta-uncertainty from the risk and ambiguity task from meta-uncertainty in the delay discounting task and decision sensitivity from both tasks. In this model, all predictors were significant (meta-uncertainty from delay discounting $\beta = 0.16, t(101) = 12.7, p < 0.001$, decision sensitivity from delay discounting $\beta = 0.04, t(101) = 2.05, p < 0.043$, and decision sensitivity from the risk and ambiguity task $\beta = -0.34, t(101) = -4.4, p < 0.001$). Model 2 predicted meta-uncertainty from the delay discounting task from meta-uncertainty in the risk and ambiguity task and decision sensitivity from both tasks. In this model, the only significant predictor was the meta-uncertainty from the other task ($\beta = 0.23, t(101) = 13.2, p < 0.001$). Model 3 was similar to model 1 but with just the decision sensitivity parameters as predictors. In this model, only the risk and ambiguity decision sensitivity parameter was significant ($\beta = -0.41, t(102) = -3.37, p = 0.001$). Model 4 mirrored model 3 for the delay discounting task meta-uncertainty. In this model neither of the decision sensitivity parameters was significant. Models 3 and 4 explained no variance in the meta-uncertainty data compared to models 1 and 2($R^2 = 0.01$ and $R^2 = -0.003$ for Models 3 and 4 versus $R^2 = 0.11$ and $R^2 = 0.49$ for Models 1 and 2). Model comparison via likelihood ratio tests confirmed that both model 1 and 2 including the meta-uncertainty parameter from the other task were better (Mod3/1 LRstat(5) = 63.4, p <0.001 and Mod4/2 LRstat(5) = 59.3, p <0.001). We conclude that the conservation across value-based tasks of metacognitive ability is not driven by correlations in decision sensitivity.
